# Determinants of Improved CGRP Peptide Binding Kinetics Revealed by Enhanced Molecular Simulations

**DOI:** 10.1101/2025.06.13.659569

**Authors:** Ceren Kilinc, Katie M. Babin, Augen A. Pioszak, Alex Dickson

## Abstract

Peptides are desirable therapeutics due to their inherent potency, safety, cost-effectiveness and ability to engage large or more complex protein surfaces. Slower kinetics of protein-peptide (un)binding can directly influence their drug efficacy and duration of action, in part by improving plasma stability of the peptide. The CLR:RAMP1 complex and its endogenous agonist peptide CGRP are of particularly high interest because of their central role in migraine pathophysiology. A better understanding of peptide binding mechanisms is needed for the development of next-generation peptide-based drugs with optimized kinetic properties. In this study, we comparatively analyze constructs of native CGRP and “ssCGRP”, an engineered variant with 430-fold longer residence time on the CLR:RAMP1 complex. Using large-scale computational resources and our high-dimensional weighted-ensemble algorithm, we then thoroughly sample and compare unbinding path ensembles for the two peptides. This elucidates the basis of the engineered residence time enhancement for ssCGRP and provides a detailed view of the intra- and intermolecular stabilizing interactions for both peptides in the bound ensemble and along the unbinding transition path. The bias-free nature of the sampling approach in combination with Markov state modeling allows for the calculation of committor values and the first analysis of protein-peptide binding transition state ensembles. Through analysis of the unbinding committor, we find that ssCGRP(27–37) also demonstrates enhanced ligand recapture of intermediate unbinding conformations and samples a more heterogeneous bound-state ensemble that entropically stabilizes the bound basin. This study shows the molecular determinants of peptide residence time at CLR:RAMP1 and provides valuable insight for the design of long-acting peptide therapeutics.

## Introduction

Calcitonin receptor-like receptor (CLR) is a class B G protein coupled receptor (GPCR) involved in various important physiological processes, including inflammatory responses^1^, vascular regulation^2^, and pain modulation^3^. The activity of CLR is regulated by the selective binding of hormones and neuropeptides, primarily consisting of three main peptide agonists: calcitonin gene-related peptide (CGRP), adrenomedullin (AM) and adrenomedullin 2/intermedin (AM2/IMD)^4^. Among these peptides, CGRP has especially attracted increasing attention due to its signaling role in chronic pain disorders and migraine pathophysiology^3,5^. CGRP is known to have protective cardiovascular effects^2,6^ and ongoing efforts have shown that it could also have functions relevant to modulating immune responses^7^, cancer^8^, and inflammatory skin conditions^9^. Given its multifunctional nature and involvement in various processes, CGRP and its receptor CLR represent critical therapeutic targets for the treatment of many disorders.

CLR is a membrane protein characterized by a common class B GPCR structure comprising a large N-terminal extracellular domain (ECD) followed by a 7 transmembrane helix domain (TMD) and a C-terminal intracellular tail. Unlike most GPCRs, which can reach the cell surface and signal independently, CLR requires the formation of a heterodimeric complex with one of the three receptor activity-modifying proteins (RAMP1–3) for proper trafficking to the cell membrane and functional signaling^10^. RAMPs are structurally composed of a large ECD followed by a single transmembrane helix and a short C-terminus. Different CLR:RAMP receptor complexes exhibit different specificities for the peptide agonists^10,11^. Specifically, CLR:RAMP1 forms the CGRP receptor, which exhibits a clear preference for CGRP over AM and AM2/IMD. In contrast, CLR:RAMP2 (AM1 receptor) and CLR:RAMP3 (AM2 receptor) favor the binding of AM and AM2/IMD, respectively, both showing a significantly lower affinity for CGRP^4^. The binding of CGRP to CLR:RAMP1 follows a two-step mechanism: initial engagement of the C-terminal ECD-binding segment of the peptide with the ECDs of the receptor complex, promoting subsequent interactions of the N-terminal TMD-binding segment of the peptide with the CLR TMD^12^. Upon peptide binding, conformational changes within the receptor predominately facilitate recruitment of heterotrimeric Gs protein, leading to intracellular cyclic AMP (cAMP) accumulation and the activation of downstream signaling pathways^13^. Figure 1a shows the stable conformation of the fully engaged CGRP peptide and the G protein trimer with the CLR:RAMP1 receptor complex.

**Figure 1:**
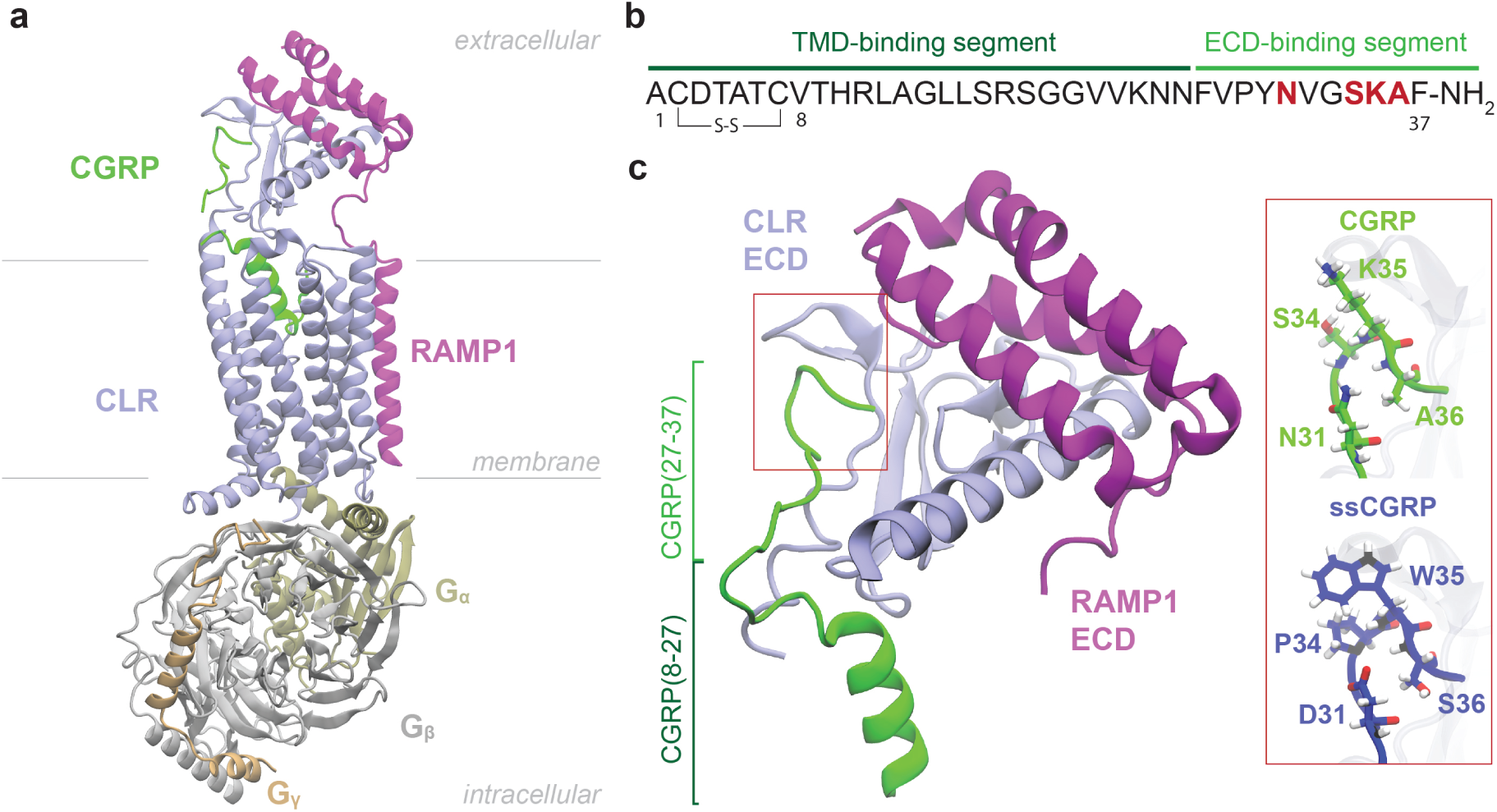
Structural overview of CLR:RAMP1 complex with CGRP peptide. (a) CGRP peptide (green) bound to CLR (iceblue):RAMP1 (purple) heterodimer complex. Binding position of Gs protein complex consisting of G*α* (tan), G*β* (gray) and G*γ* (beige) and orientations of receptors in the membrane are also shown. Structure from PDB ID 6E3Y^14^.(b) Top: CGRP primary sequence. The sequences of two binding segments are color coded. The mutated residues are marked in red. Bottom: The ECD part of the receptors bound with CGRP and CGRP’s sequences of two binding segments on its structure are shown. Structure from PDB ID 4RWG^15^. The four mutated residues in the ssCGRP peptide are highlighted in the red box.

The two-domain binding model highlights the respective roles of ECD and TMD in CGRP binding, where the ECD-binding segment of CGRP binds to the ECD with high affinity, anchoring the ligand and enabling initial receptor recognition, while the TMD-binding segment binds to the TMD with lower affinity, a step essential for full receptor activation (Fig. 1b). Cryo-EM structures have shown that in the absence of the Gs protein, the CGRP agonist predominantly engages ECD, with little to no stable interaction with TMD^16^, while in the presence of the Gs protein, the CGRP agonist binds to both domains^14^. In addition, full antagonist activity was observed using short peptide fragments that bind only to the receptor ECD, indicating that preventing the initial ECD-mediated binding step is sufficient to effectively antagonize the receptor^17^. Therefore, the ECD of the CLR:RAMP1 complex provides extensive and valuable opportunities for selective targeting by small molecules, peptides, and antibodies. Several small molecule^18–23^ and monoclonal antibody^24–30^ therapeutics, which bind to the ECD of the CGRP receptor^31–33^ or the ECD-binding segment of CGRP^34^, have been approved by the US Food and Drug Administration (FDA) for acute and preventive migraine treatment. Despite the success of small molecules and antibodies in targeting the CGRP receptor, peptides remain a desirable therapeutic alternative due to their inherent potency, safety, cost-effectiveness and ability to engage larger or more complex protein surfaces; however, they often suffer from poor plasma stability and rapid clearance^35^. Recent efforts have focused on modifying the pharmacokinetic profiles of peptide antagonists for CGRP to improve their metabolic stability and prolong their residence time on the receptor^36–40^. However, no CGRP peptide antagonist has yet progressed into clinical trials as a therapeutic candidate. This could be due in part to a significant gap in our understanding of peptide–receptor interactions at the molecular level. Most current models of peptide binding, including for FDA-approved drugs, are based on static crystal structures or short-timescale molecular dynamics (MD) simulations. Although these approaches have identified important binding contacts, they offer limited understanding of the evolution and persistency of important interactions during the critical ligand unbinding process, which is particularly relevant for the engineering of long residence-time peptides. In particular, the dissociation mechanism of native CGRP and how it differs from that of a long residence-time peptide remain unexplored.

To fill this critical gap, this study presents the first, to the best of our knowledge, comparative mechanistic analysis of peptide ligand dissociation from a protein target. The study focuses on exploring the unbinding pathways and kinetics of the native CGRP antagonist and an ultrahigh-affinity CGRP variant designed by the Pioszak lab, with four residue substitutions near the C-terminus of CGRP (Fig. 1c). The “ssCGRP” variant, named after its sustained cAMP signaling in its full-length agonist form, binds the ECD of the CGRP receptor with ≈1000-fold higher affinity and prolonged residence time, while also showing kinetic selectivity for the CGRP receptor complex over other RAMP pairings^41,42^. Long residence time antagonists of the ssCGRP variant were generated by N-terminal truncation. Exploration of the determinants of these prolonged kinetics requires methods that can explore steep energy landscapes, broadly sample different transition paths, and compute unbiased transition rates. This is a great challenge due to the long timescales associated with peptide unbinding paths (milliseconds to minutes), which are roughly a million-fold beyond the reach of conventional molecular dynamics (MD) simulations. Weighted ensemble (WE) is a powerful enhanced sampling method in which an ensemble of trajectories with statistical weights is propagated forward in time together and adaptively resampled by merging and cloning to efficiently sample long-timescale events^43,44^. WE introduces no bias potentials and thus preserves the underlying free energy landscape, whereas bias-driven enhanced sampling methods apply external forces and include some approximations that can locally reshape the energy surface and obscure the authentic ensemble of transition paths^45^. Additionally, peptide ligands exhibit extensive conformational flexibility compared to small-molecules, requiring extensive sampling of their transition paths, and posing problems for enhanced sampling algorithms that employ biasing forces along one- or two-dimensional reaction coordinates.

In this study, we leverage recent advances in path sampling methods by combining the WE-based REVO (resampling of ensembles by variation optimization) method^46^ with Markov state models (MSM)^47,48^, which approximate long-timescale kinetics by putting together many short simulations into a network of transitions between discrete conformational states. The ability of this approach to capture extensive ensembles of unbinding pathways for long-timescale processes (up to minutes in duration) has been shown for small molecule unbinding across various protein targets^46,49–51^. Here, we show that this can be extended to sample unbinding trajectories of flexible peptides and compute their unbiased kinetics. Using this framework, we simulate over 100 peptide unbinding events in order to decipher the molecular determinants of the prolonged ssCGRP residence time, while obtaining qualitatively consistent unbinding kinetics with biolayer interferometry (BLI) experiments.

## Results

### Experimental Determination of CGRP and ssCGRP Antagonist (8-37) Binding Kinetics

The CLR:RAMP1:CGRP complex represents a highly dynamic and flexible receptor-peptide system, which poses significant challenges for structural and kinetic characterization. To experimentally determine the binding kinetics of CGRP and ssCGRP to the CGRP receptor ECD, we performed biolayer interferometry (BLI) using a purified CLR:RAMP1 ECD fusion protein expressed in HEK293T cells. The soluble CLR:RAMP1 ECD complex was engineered as a fusion of the two ECDs connected by a flexible linker and with a C-terminal His-6 tag similar to our previous studies^52,53^. The fusion protein was secreted from HEK293T cells and purified by nickel affinity and size exclusion chromatography (Supplementary Fig. 1). The fusion protein was eluted from the size exclusion column as a symmetric peak. Treatment of the pooled sample with PNGase F yielded a single band around 25 kDa, indicating that the recombinant CLR:RAMP1 ECD fusion protein was N-glycosylated as expected.

Biolayer interferometry (BLI) was used to measure the kinetics of the interaction of the fusion protein with wildtype and ssCGRP peptides immobilized on streptavidin biosensor tips. N-terminally biotinylated (8-37) antagonist fragments of CGRP and ssCGRP were used to provide an extended distance between the immobilization site and the (27-37) portion of the peptides that binds the ECD complex. The association and dissociation phase data for a series of ECD complex concentrations were globally fit to an association and dissociation equation to determine the kinetic parameters. Purified CLR:RAMP1 ECD complex binding to Biotin-CGRP(8-37) resulted in a *k*_on_ of 6.33 × 10^4^ M^−1^s^−1^, a *k*_off_ of 3.3 s^−1^, and a calculated affinity (*K_D_*) of 52.2 *µ*M (Fig. 2a,c,d, Supplementary Table 1).

**Figure 2:**
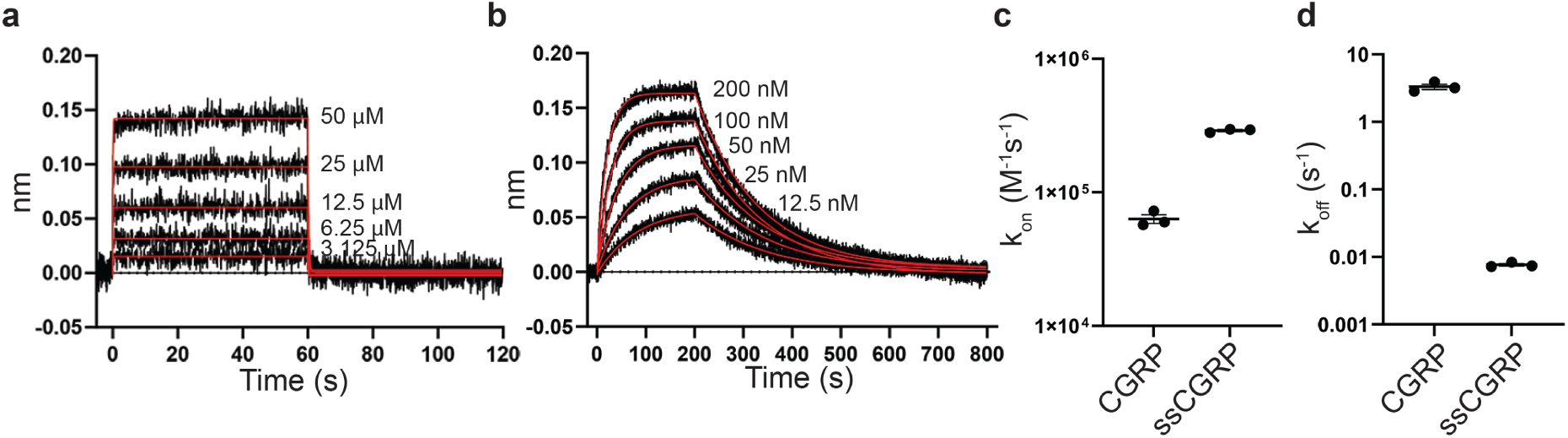
Association and dissociation of CGRP and ssCGRP at purified CLR:RAMP1 ECD complex using BLI. (a) BLI traces of immobilized biotin-CGRP(8-37) on streptavidin biosensor tips binding and then unbinding from purified CLR:RAMP1 ECD complex. Data was collected at 10 Hz. Kinetics curve fits are shown in red. (b) Similar data to (a), but with biotin-ssCGRP(8-37). Data was collected at 5 Hz. (c,d) Scatter plot showing the three independent replicates ± SEM of the association rates (c) and dissociation rates (d).

This yielded a residence time of 0.3 seconds and a half-life of 0.2 seconds. In contrast, binding of the ECD complex to biotin-ssCGRP(8-37) showed much slower association and dissociation phases. These yielded an on rate of 2.90 × 10^5^ M^−1^ min^−1^, an off rate of 0.0077 s^−1^, and a calculated affinity of 26.4 nM, resulting in a residence time and half-life of 2.18 minutes and 1.52 minutes, respectively (Fig. 2b-d, Supplementary Table 1). These results indicated that the purified CLR:RAMP1 ECD complex had ∼2,000-fold stronger affinity and a 4.5-fold faster on rate and 430-fold slower off rate for the ssCGRP peptide as compared to the wt peptide. The findings of the BLI assay clearly demonstrate the enhanced stability of the ssCGRP peptide and provide a solid means of connection with unbinding rates determined computationally through enhanced weighted ensemble sampling.

### Unbinding Simulations of CGRP and ssCGRP Antagonists (8-37)

To investigate why ssCGRP(8-37) has a slower disassociation rate than CGRP(8-37) at the molecular level, we initially constructed two simulation systems designed to match the experimental setup used in the BLI assays. Each system contains the ECDs of the receptor proteins, comprising residues 29–134 of CLR and 24–111 of RAMP1, in complex with either CGRP(8–37) or ssCGRP(8–37). For each system, we performed 6 runs of unbinding simulations that start from an equilibrated crystallographic bound pose and use WE simulations with the REVO algorithm. Each run has at least 7500 cycles and a total cumulative simulation time of 87.84 *µs* was reached across all (8-37) runs (Table 1).

**Table 1:** Summary of REVO simulation parameters and unbinding statistics for each system. The unbinding distance is calculated using two reaction coordinates: the RMSD of the entire peptide (“all”) or the RMSD of the C-terminal fragment (32-37) of the peptide (“tail”). The cumulative simulation time is calculated by summing the lengths of all of the 48 walkers in each set of simulations. The num. of unbinding events indicated how many independent peptide disassociation were observed in each set of run and is determined by counting each walker reaching to a minimum peptide-receptor interatomic distance greater than 10.0 Å. The table also lists the total unbinding weights and the estimated mean first passage time for each run.

| System | Distance selection | Num. of runs | Cycles per run | Cum. simulation time ($\mu\text{s}$) | Num. of unbinding | Total unb. weights | MFPT (s) |
| --- | --- | --- | --- | --- | --- | --- | --- |
| CGRP(8-37) | all | 3 | 7750 | 22.32 | 1 | $7.0 \times 10^{-6}$ | $6.7 \times 10^{-2}$ |
|  | tail | 3 | 7500 | 21.6 | 0 | 0 | 0 |
| ssCGRP(8-37) | all | 3 | 7750 | 22.32 | 0 | 0 | 0 |
|  | tail | 3 | 7500 | 21.6 | 0 | 0 | 0 |
| CGRP(27-37) | all | 6 | 5500 | 31.68 | 18 | $6.64 \times 10^{-2}$ | $9.95 \times 10^{-6}$ |
| | tail | 6 | 5500 | 31.68 | 71 | $1.0 \times 10^{-2}$ | $6.62 \times 10^{-6}$ |
| ssCGRP(27-37) | all | 6 | 5500 | 31.68 | 3 | $3.43 \times 10^{-6}$ | $1.92 \times 10^{-1}$ |
| | tail | 6 | 5500 | 31.68 | 14 | $2.90 \times 10^{-11}$ | $2.27 \times 10^4$ |

As mentioned in Methods, two definitions of the inter-trajectory distance were used in our REVO calculations: one that measures conformational differences in the peptide C-terminal tail spanning from residues 32 to 37 (“tail”) and one that measures differences in the entire peptide (“all”). This only affects how trajectories in the weighted ensemble algorithm are merged or cloned, and thus only affect the efficiency of sampling and not the long-time behavior of the results. All simulations are run in the “unbinding ensemble”^46,49^ where trajectories begin in the peptide-bound state and are terminated in the unbound state, as defined by achieving a minimum peptide-receptor interatomic distance greater than 10.0 Å. Only one such unbinding event occurred for the 8-37 simulation set, which was for the CGRP peptide.

The weights of the walkers are used to calculate all observables from the set of simulations. A non-equilibrium “free energy” (NEFE) profile can be computed using a weighted histogram along a set of bins: *W_j_* = −ln ∑*_i_*_∈_ *_j_ p_i_*, where *p_i_* is the weight of the *i*^th^ frame and *W_j_* is the “free energy” of the *j*^th^ bin. We note that this is not equivalent to an equilibrium free energy, as trajectories that originate from the unbound state are not included. The NEFE projected along the tail RMSD is shown in Fig. 3a. Due to their conserved and relatively more stable beta-turn structure, the *Cα* atoms of the last 6 residues (32-37) in the C-terminal tail were used to calculate the RMSD of the peptide from the bound state after alignment to the CLR and RAMP1 structures in the starting pose. Throughout the manuscript, all references to RMSD specifically denote the tail RMSD. The NEFE is calculated using 20 bins and each curve is vertically shifted so that its minimum is zero.

**Figure 3:**
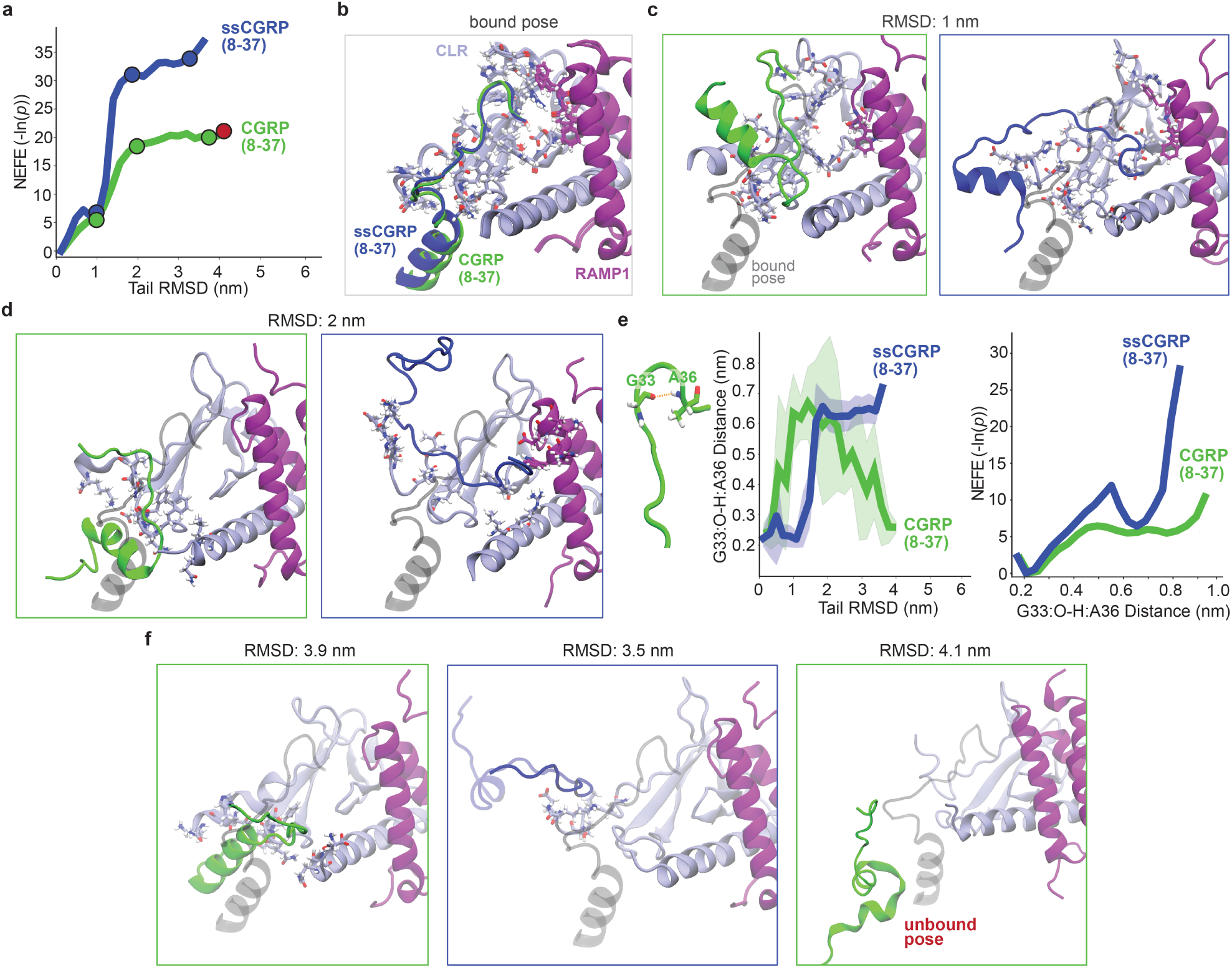
Structural characteristics of peptide 8-37 segment unbinding obtained from REVO simulations. (a) Non-equilibrium free energy (NEFE) profiles of CGRP(8-37) (green) and ssCGRP(8-37) (blue) along the tail (32-37) RMSD of peptides. The RMSD points where conformation snapshots are chosen are shown in red for the CGRP unbound conformation. (b,c,d) Representative snapshots corresponding to tail conformations at bound pose (b), 1.0 nm (c) and 2.0 nm (d) for CGRP(8-37) (green) and ssCGRP(8-37) (blue). The starting peptide bound pose is shown in gray. CLR (ice blue), RAMP1 (purple) and residues within 0.3 nm distance to peptides are included in all snapshots. (e) (left) Evolution of the distance between the main oxygen atom of G33 and the hydrogen atom in the main nitrogen of A36 as a function of tail RMSD and (right) NEFE profiles along this distance are shown for both peptides. The shaded areas show the standard deviation of distances. The corresponding residues is shown on the ligand structure at the left. (f) Representative snapshots corresponding to tail conformations at near-unbound poses for CGRP(8-37) (left) and ssCGRP(8-37) (middle), and the CGRP(8-37) unbound pose (right).

The plots combine data from all WE runs to provide insight into the relative stability of metastable states and reveal the energetic barriers governing ligand dissociation. As the RMSD increases, indicating a greater deviation from the initial bound conformation, the free energy increases for both peptides. Notably, the ssCGRP(8-37) profile displays a steeper rise and a consistently higher free energy across the reaction coordinate compared to CGRP(8-37). Both peptides experience a barrier of ≈ 5-7 kT prior to achieving a RMSD of 1.0 nm. A dramatic difference in stability between the peptides is then observed as they surpass a secondary barrier towards an RMSD of 2.0 nm. The conformation snapshots of the trajectories at 1 nm (Fig. 3c) and 2 nm (Fig. 3d) RMSDs showed that the stable beta-turn structure between G33 and A36 was distorted at a higher RMSD value compared to its initial bound pose (Fig. 3b).This is quantified by calculating the beta turn distance between the main oxygen atom of G33 and the hydrogen atom in the main nitrogen of A36 (Fig. 3e). The beta-turn distance is estimated using weighted histograms with 20 bins for all frames, the same manner as the NEFE profile, and plotted as a function of the tail RMSD from the bound state for CGRP(8–37) and ssCGRP(8–37). This distance captures the formation and disruption of a characteristic beta-turn motif within the peptide during unbinding. For both peptides, the beta-turn distance increases sharply between RMSD values of 1.0 and 2.0 nm, and this happens at lower RMSDs for CGRP. At higher RMSDs the beta-turn distance for CGRP again decreases, although this is highly variable and based on only a few high-RMSD trajectories. The corresponding NEFE profile of the beta-turn distance further supports the enhanced stability of ssCGRP, which again has a higher energetic barrier to dissociated beta-turn state. These structural trends correspond well to the free energy profiles shown in panel 3a. For both peptides, the 1–2 nm RMSD region corresponds to a critical transition zone where breaking interactions at the receptor interface and the destabilization of the secondary structure together significantly increase the energetic barrier to further dissociation. These findings show that ssCGRP faces higher energetic barriers during unbinding, consistent with its slower off-rate and prolonged residence time observed experimentally.

When there is sufficient sampling of unbinding events, WE simulations can be used to directly estimate mean first passage times (MFPTs), which is defined as the average time required for a ligand to reach the unbound state from the bound state (Eq. 5) and is the inverse of the residence time. Although some trajectories reached near-unbound conformations for CGRP(8-37) (RMSD:3.9 nm) and ssCGRP(8-37) (RMSD:3.5 nm), the peptide does not fully disengage from the receptor in any case for ssCGRP(8-37) and only once for CGRP(8-37) (RMSD:4.1 nm) (Fig. 3f). This suggests a stronger binding to the receptor and substantially longer residence time for ssCGRP, which is consistent with experimental measurements. For CGRP(8-37), the MFPT is estimated as 0.13 s using the unbinding weight of 7.0 × 10^−6^, which can be directly compared with the CGRP(8-37) MFPT obtained from the BLI assay of 0.3 s. The computational MFPT is ≈ 2.3 times faster than the experimental MFPT but still within the same order of magnitude. However, sampling for the 8-37 systems is not enough to support a robust analysis of the underlying unbinding mechanisms. Both CGRP(8-37) and ssCGRP(8-37) are relatively long and highly flexible peptides, allowing them to adopt extended conformations that introduce a high degree of structural variability and sampling complexity. To mitigate this, we focus next on a truncated version of the peptides that includes only the ECD-binding segment, which is known to initially engage with the receptor and still contains all of the sequence differences between CGRP and ssCGRP.

### Unbinding Simulations of CGRP and ssCGRP ECD-binding segments (27-37)

We then performed unbinding simulations for systems containing the shorter ECD-binding segments of CGRP(27–37) or ssCGRP(27–37) in complex with CLR:RAMP1 ECD. These are initialized from an equilibrated bound pose using WE-REVO simulations. Each of the 12 runs per system has 5500 cycles, and the total cumulative simulation time across all runs is 126.72 *µ*s (Table 1). The NEFE profiles projected along the ligand tail RMSD are shown in Fig. 4a. ssCGRP(27-37) exhibits a higher free energy increase compared to CGRP(27-37) throughout and shows a substantially steeper free energy rise starting at RMSD values around 4 nm. It is worth to keep in mind that the average RMSD value for unbound states in CGRP(27-37) is 3.57 nm and in ssCGRP(27-37) is 3.41 nm (red circles in Fig. 4a). Thus, regions in the RMSD curves beyond the red circles correspond to rare unbinding events in which the peptide achieves a high distance from the binding site while still making transient contact with the ECD. These conformations are highly committed to unbinding and are not relevant for the determination of ligand binding kinetics or free energies. Both peptides experience a gradual increase in free energy between ≈ 5 - 15 kT between 1.0 - 3.0 nm. This increase in free energy is not as sharp as it is for 8-37 systems, indicating a higher probability of unbinding and a more gradual unbinding process.

**Figure 4:**
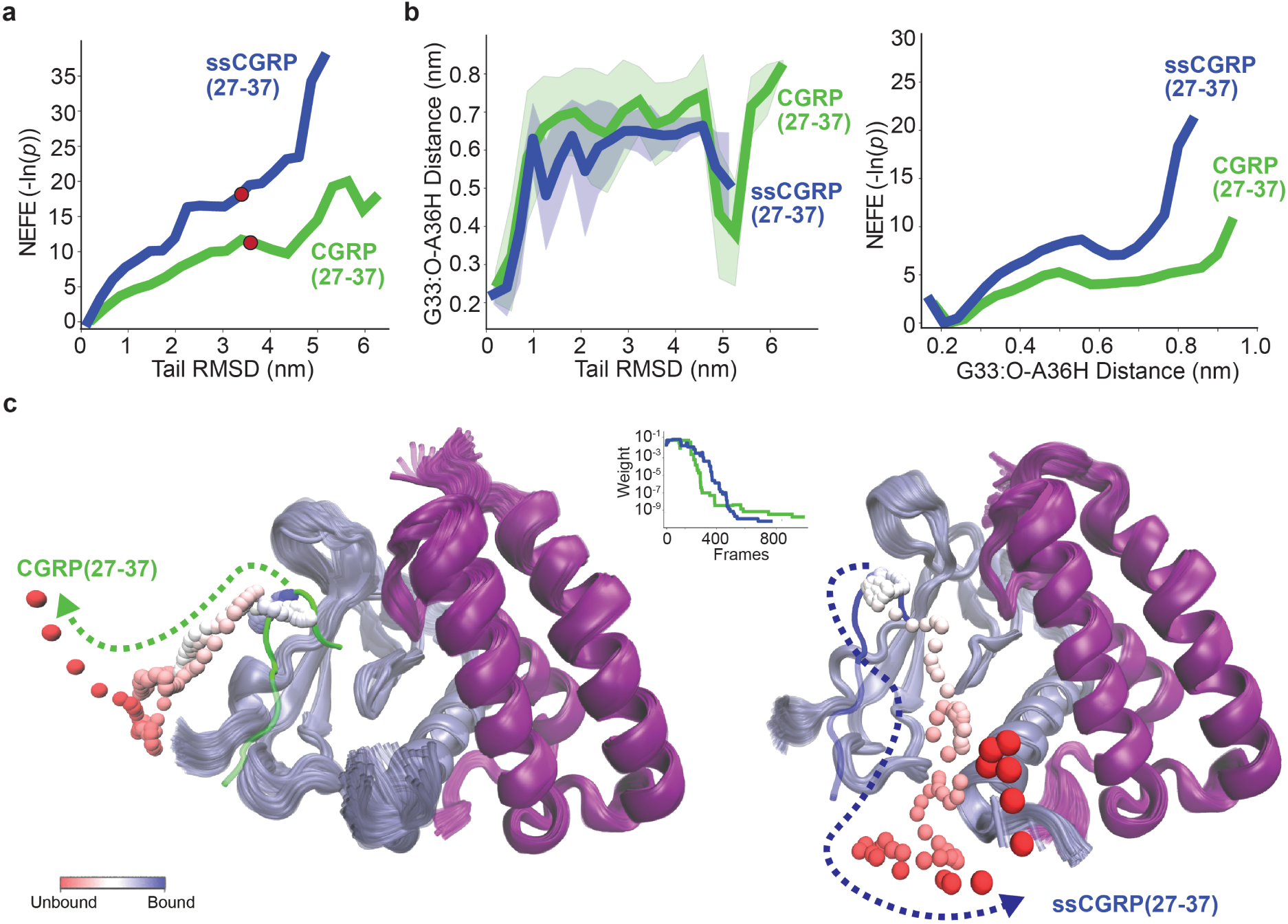
Energetic and structural characteristics of peptide (27-37) segment unbinding obtained from REVO simulations.(a) NEFE profiles of CGRP(27-37) (green) and ssCGRP(27-37) (blue) along the tail (32-37) RMSD of peptides. The RMSD points where conformation snapshots are chosen are shown in corresponding colors to the peptide for the rest and mean RMSD value for the unbound conformations shown in red. (b) (left) Evolution of the beta-turn distance between the main oxygen atom of G33 and the hydrogen atom in the main nitrogen of A36 as a function of ligand tail RMSD for both peptides computed from REVO simulations. The shaded areas show the standard deviation of distances. (right) The NEFE profiles along beta-turn distance are shown for both peptides. (c) The representative unbinding pathway trajectories from REVO simulations for CGRP(27-37) (left) and ssCGRP(27-37) (right). The residue 34 on peptides is selected as middle point of the tail region (shown as ball representation) to show the unbinding pathways along the trajectory and is color-mapped from blue (bound) to red (unbound) to reflect progression along the unbinding coordinate. The weight change along the trajectories are shown in the graph for both peptides.

In contrast to the 8-37 systems, both 27-37 peptides now display a rapid increase in the beta-turn distance as RMSD grows to ≈ 1.1 nm (Fig. 4b), marking the initial disruption of the beta-turn secondary structure at the beginning of unbinding. At higher RMSDs, both peptides exhibit a greater fluctuation in the beta-turn distance with a high standard deviation, suggesting significant structural flexibility in the dissociated state. This NEFE profile of the beta-turn distance is consistent with the RMSD profiles, where the ssCGRP free energy rises more steeply with distance, indicating that breaking this internal contact along the dissociation pathway is energetically more unfavorable. ssCGRP(27-37) again exhibits a significantly steeper increase in free energy compared to CGRP(27-37), although the 27-37 system free energy profiles are generally lower than those for the 8-37 systems. This indicates that the 8-27 regions of both peptides help anchor the peptide-CLR interactions and increase energetic barriers along the unbinding pathway, particularly in the RMSD region of 1-2 nm. It is unclear whether the differences in the behavior of the ssCGRP beta-turn are due to better sampling in the 27-37 systems, or are inherent to the peptide structure. Regardless, we consistently observe that deformation of the beta-turn is required for peptide unbinding and that this deformation is less favorable for ssCGRP.

A total of 89 and 17 unbinding events were observed for CGRP(27-37) and ssCGRP(27-37), respectively, summed over all weighted ensemble simulations. Representative unbinding pathways are shown in Fig. 4c and Supplementary Video 1-Supplementary Video 2. For CGRP(27-37), the MFPT is calculated as 7.9 × 10^−6^ s, with a total accumulated trajectory weight of 1.7 × 10^−1^. In contrast, ssCGRP showed slower unbinding with an MFPT of 3.8 × 10^−1^ s and a weight of 3.4 × 10^−6^. Naturally, 27-37 systems showed faster disassociation than 8-37 systems for both peptides. It is important to note that in our direct WE rate estimates, a relatively large uncertainty arises because of the limited sampling of unbinding events. In this approach, rate calculations depend only on the statistical weights of the reactive trajectories that reach the unbound state. Previous work has shown that unbinding rates calculated through the construction of a Markov state model (MSM) show less variability as they incorporate the full set of simulation data^49^.

#### Unbinding Pathways and Kinetics from Markov State Modeling

MSM-derived rates are less sensitive to the limited sampling of rare dissociation events as they capture contributions from intermediate or partially dissociated conformations, even if those trajectories do not complete the unbinding process within the simulation timescale. To provide insight into mechanisms, committors, and transition rates, we constructed separate MSMs for each of the CGRP(27-37) and ssCGRP(27-37) systems using all data obtained from REVO simulations. The conformational space was first featurized using distances between C*α* atoms on the peptide and on the binding site of the receptor complex. This is followed by reduction of dimensionality using time-lagged independent component analysis (tICA)**^?^**and clustering of these data into discrete microstates using the KMeans algorithm (see the Markov State Modeling section in Methods for more detail). We systematically varied the number of tICA output dimensions (3, 5, 8, and 10) and the number of clusters (50, 200, 500, 800, 1000) used for discretization (Supplementary Fig. 2) and built 5 MSMs for each parameter set. The MSM-derived MFPTs for CGRP remained consistent with the flux-based estimates in all combinations of parameters, with values ranging around 10^−6^ s for CGRP(27-37), while the values for ssCGRP(27-37) showed greater variability, with MFPTs from individual MSMs ranging from 10^−5^ s to 10^−1^ s. For all parameter values tested, the mean MFPTs for ssCGRP remained consistently around 0.03 s, substantially slower than those of CGRP. The trends we observed for residence times are qualitatively consistent with experimental results, and the MFPTs derived from these MSMs demonstrated robustness, varying by less than an order of magnitude between different combinations of parameters. The final transition probability matrices used for the analysis of each system were constructed with 3 tICA output dimensions and 1000 states. These have mean MFPTs of 7.5 × 10^−6^ s and 0.02 s for CGRP(27-37) and ssCGRP(27-37), respectively.

Transition probability matrices from MSM construction were used to generate conformation space networks (CSNs) of CGRP(27-37) and ssCGRP(27-37). In Figure 5a (top), each CSN has 1000 nodes where each node corresponds to a particular cluster of peptide–receptor conformations, the size of the nodes corresponds to the sum of the WE weights of these conformations, and the edges represent transitions between nodes were observed in a single WE cycle. Representative peptide–receptor conformations for each cluster can be found in Supplementary Trajectory 1 and Supplementary Trajectory 2. Each network is color-coded according to the tail RMSD or the unbinding committor probability (*p_B_*_→*U*_). When colored by tail RMSD, both peptides show a general progression from low-RMSD bound states (dark blue) to high-RMSD unbound states (red). CGRP(27-37) shows a relatively compact and densely populated low-RMSD region, suggesting reduced stability in the bound state. In contrast, ssCGRP(27-37) samples a broader distribution of bound states and shows a larger separation between the bound and unbound regions of the network, indicating a slower, more gradual transition toward unbinding. The networks are also colored by the unbinding committor probability, which quantifies the likelihood that a system initialized at an intermediate state will transition (commit) to the unbound basin before the bound basin. For states lying outside the defined bound and unbound basins, the committor spans values between 0 (likely to rebind) and 1 (likely to unbind). In Figure 5a (middle), these probabilities are mapped to a color gradient, with blue corresponding to the bound states (*p* = 0), red to the unbound states (*p* = 1), and the transition state ensemble (TSE) shown in yellow. The CGRP network shows a relatively short jump from low to high committor probability states, indicating a sharper transition to unbinding and a clear distinction bound-committed and unbound-committed conformations. In contrast, ssCGRP exhibits a more continuous and distributed progression across the network.

**Figure 5:**
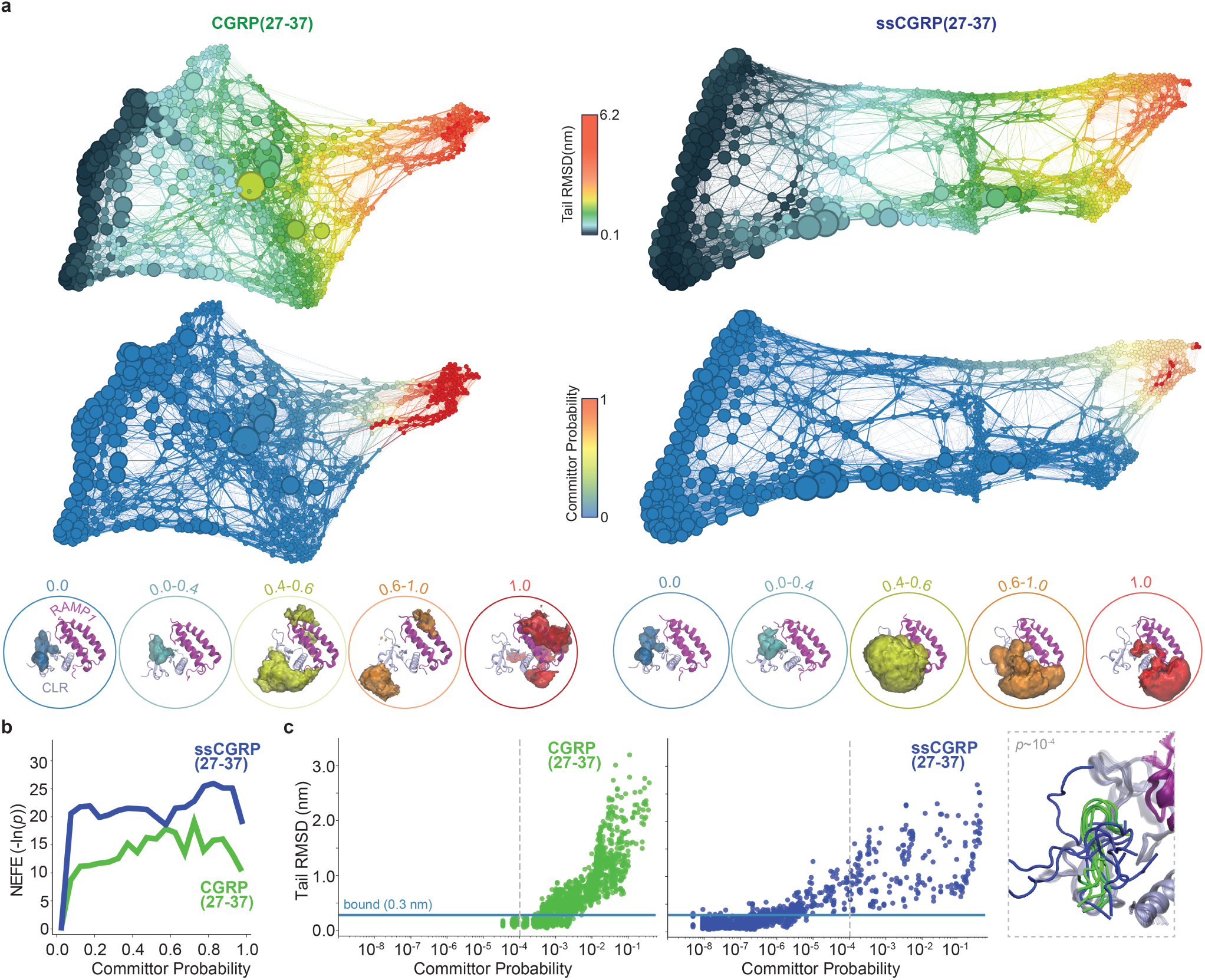
Conformational space network of CGRP(27-37) and ssCGRP(27-37) unbinding from the CLR:RAMP1 ECD complex. (a) CSN for CGRP(27-37) (left) and ssCGRP(27-37) (right), colored by the tail RMSD of the ligands (top row) and unbinding committor probability (middle row). The bottom row shows the density volume maps of spatial distributions of peptide conformations projected onto the receptor complex (CLR in ice blue, RAMP1 in purple), computed across committor probability bins: 0.0 (blue), 0.0–0.4 (cyan), 0.4–0.6 (yellow), 0.6–1.0 (orange), and 1.0 (red). Density surfaces were rendered in VMD using an isovalue of 0.05 for the first two bins and 0.005 for the remaining bins. (b) NEFE profiles as a function of committor probability for CGRP(27-37) (green) and ssCGRP(27-37) (blue). (c) The tail RMSD of the CGRP(27-37) (left, green) and ssCGRP(27-37) (right, blue) plotted as a function of committor probability on a log-scale. The horizontal blue line marks the 0.3 nm bound-state cutoff, and the vertical dashed line indicates the committor value at 10^−4^. Representative ligand conformations corresponding to this probability are shown to the right in the dashed gray box.

For more systematic determination of structural changes along the unbinding pathways, we grouped all data based on committor probability values where the data set was discretized into five distinct committor bins: the bound bin (*p* = 0), the near-bound bin (0 *< p <* 0.4), the transition bin (0.4 ≤ *p* ≤ 0.6), the near-unbound bin (0.6 *< p <* 1), and the unbound bin (*p* = 1). This provides a coarse-grained view of the progression from bound to fully unbound states along the unbinding committor. For the CGRP(27-37) system, the number of clusters in each bin was: 12 (bound), 852 (near-bound), 11 (transition), 24 (near-unbound), and 101 (unbound) and the number of conformations in each bin was: 149776, 2901984, 7555, 13043, and 95690, respectively. For the ssCGRP(27-37) system, we identified 34 (bound), 821 (near-bound), 94 (transition), 43 (near-unbound), and 8 (unbound) clusters with number of 759269, 2360948, 33895, 10979 and 2957 conformations, respectively. Both CGRP(27-37) and ssCGRP(27-37) exhibit extensive sampling within the bound and near-bound committor bins, but differ in their exploration of the transition and unbound states. ssCGRP displays broader sampling across the transition and near-unbound regions but shows a steep decline in occupancy within the fully unbound bin. In contrast, CGRP shows relatively sparse sampling of the transition state ensemble, with a gradual increase in representation toward the unbound state. The density volume maps of each bin using 3000 randomly selected frames are constructed to reveal the spatial distribution explored by the ligands along the committor probability. Non-weighted density maps are provided in Fig. 5a (bottom) as isosurfaces for each bin. Density maps form a highly compact region for the bound and near-bound bins located within the receptor binding pocket on the CLR. In contrast, the transition, near-unbound and unbound bins contain more diffuse density maps covering regions on both CLR and RAMP1.

Figure 5b shows the NEFE profiles for both peptides as a function of the unbinding committor probability. While CGRP(27-37) maintains a relatively lower free energy profile across all committor probabilities, both peptides exhibit a sharper increase in free energy at the initial stages of unbinding, when compared with NEFE profiles of the tail RMSD. In this transition, the free energy increases to ≈ 10 kT for CGRP(27–37) and ≈ 20 kT for ssCGRP(27–37), which constitutes roughly the entire free energy cost of unbinding for both peptides. As expected, this is in accordance with the NEFE unbinding free energy as a function of tail RMSD, using the red circles to indicate the unbound state in Fig. 4a.

To investigate the earliest stages of the unbinding process, we examine the ligand tail RMSD as a function of the committor plotted on a log scale (Fig. 5c). For CGRP(27-37), a steep transition is observed starting near a committor of 10^−4^ with RMSD values increase rapidly above the bound state cutoff of 0.3 nm. Using a committor value of 10^−4^ as a guide (dashed line), we see a much broader distribution of RMSD values for ssCGRP than CGRP. The presence of high-RMSD, low-committor conformations in ssCGRP reflects a greater degree of “ligand recapture”, where movement away from the bound basin is less correlated with unbinding progress. For ssCGRP, the 10^−4^ committor structures with a tail RMSD *>* 1.0 nm have lost most if not all of their native contacts (Fig. 5c (right)), yet still overwhelmingly commit to the bound state (9999 out of 10000 times).

### Interaction Network of CGRP(27-37) and ssCGRP(27-37)

Figure 6 provides a detailed comparison of the residue-level interaction patterns that differentiate the binding behavior of CGRP(27–37) and ssCGRP(27–37) to the CLR:RAMP1 ECD. Contacts between ligand and receptor residues were defined using a minimum distance cutoff of 2.5 Å between the nearest atoms. Contact frequencies were computed for each frame and averaged within each committor bin to capture ensemble-level interactions along the unbinding pathway. Only residue pairs with an average contact probability exceeding 5% are shown in the heatmaps (Fig. 6a) to emphasize the most common interactions. Contact frequency maps show that, although both peptides mainly engage the same regions of the receptor, they differ in the persistence of those contacts. We construct contact-difference maps by taking, for each residue, the frame-wise contact count for ssCGRP minus that for CGRP. In these plots, positive values indicate residues more frequently contacted by ssCGRP (blue), whereas negative values highlight those preferentially engaged by CGRP (green). In the bound state, ssCGRP shows enhanced contacts with the CLR residues H117, N118, R119, and W121, whereas CGRP primarily engages H114 and R119. In particular, W35^ssCGRP^ has persistent interactions with H117, N118, and R119 incorporating all turret loop residues, and the corresponding K35^CGRP^ engages only with R119 from this region. Although ssCGRP recapitulates every contact formed by CGRP in this region, S34^CGRP^ shows especially strong persistence in H114 and R119. In RAMP1, the *α*2 − *α*3 loop residue F83 forms a persistent contact with W35^ssCGRP^, whereas this residue on CGRP instead engages K35^CGRP^ and F37^CGRP^. F37^CGRP^ has a wide interaction range, contacting the RAMP1 residues W74, F83, W84, and P85, while the corresponding F37^ssCGRP^ interaction is restricted to only W84 and P85. In the near-bound state, ssCGRP begins to create more persistent interactions on the hydrophobic patch below the CLR turret loop than those of CGRP in addition to the aforementioned interactions in the bound state. In particular, V32^ssCGRP^ interacts with W121, Y124, and T125 with a higher frequency, while this residue in CGRP shifts its interaction to residue F95 in the *β* 3 − *β* 4 loop. *β* 3 − *β* 4 loop residues D94 and F95 further support the interaction with T30^ssCGRP^. This shift also affects the N-terminus of CGRP(27-37) and ssCGRP(27-37) where F27^CGRP^ and V28^CGRP^ interact more with the *β* 3 − *β* 4 loop residues and these interactions stay weak for F27^ssCGRP^ and V28^ssCGRP^. In RAMP1, the *α*2 − *α*3 loop residue F83 protects its interaction with W35^ssCGRP^ and F37^CGRP^ while the rest of the interactions between the peptide and RAMP1 start to fade for both peptides. The crucial differences in interaction patterns are highlighted with the orange boxes in the panel, and these key distinctions are shown in the CLR:RAMP1 structure in Fig. 6b.

**Figure 6:**
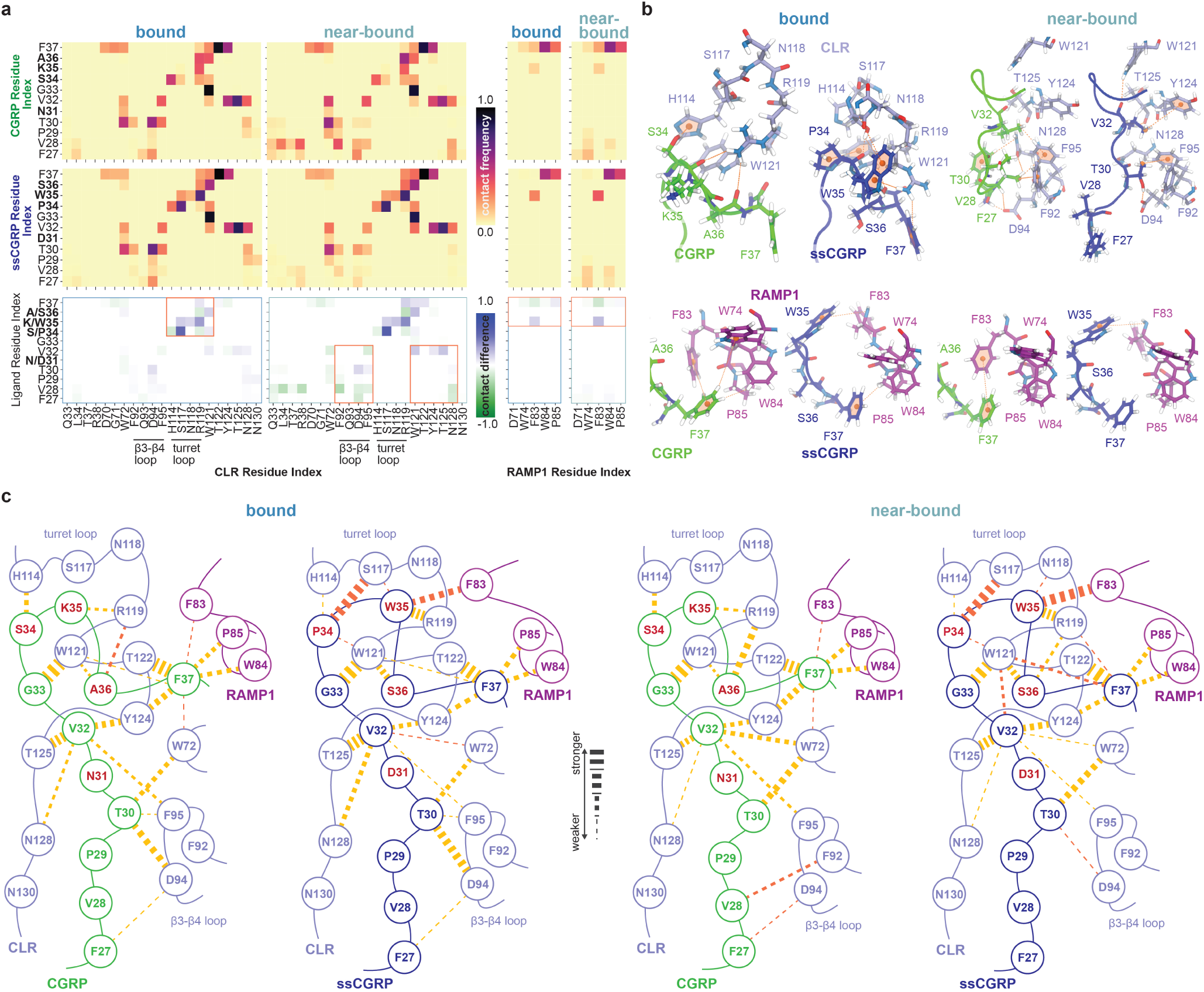
Residue-level interaction analysis for CGRP(27–37) and ssCGRP(27–37) with the CLR:RAMP1 ECD receptor complex along their unbinding pathways. (a) Heatmaps of contact frequencies (top and middle rows) and their differences (bottom row) between peptide residues and either CLR (left panels) or RAMP1 (right panels) for the bound and near-bound ensembles. Warmer colors indicate more frequent contacts, while cooler tones indicate lower contact frequency. Contact difference maps use a diverging color scale to reflect the enrichment of specific interactions in CGRP (negative, green) or ssCGRP (positive, blue). Regions selected for further structural analysis in panel (b) are boxed in orange. (b) Representative structural snapshots from bound and near-bound bins depict key interaction interfaces for CGRP (green) and ssCGRP (blue) with CLR (ice blue, top) and RAMP1 (purple, bottom). Key peptide-receptor contacts are shown with dashed orange line and hydrophobic cores on the residues are marked with ball representation. (c) Schematic residue interaction networks for CGRP and ssCGRP illustrate differences in contacts with CLR and RAMP1. Key peptide-receptor residue contacts with frequencies above 0.2 are shown as dashed orange lines, with line thickness proportional to contact frequency (thicker lines indicate stronger interactions). Contacts unique to a specific peptide system are highlighted in dark-orange.

Figure 6c illustrates the extensive contact network between the peptides and the CLR:RAMP1 receptor complex in the bound and near-bound states for interactions with more than 0.2 contact frequency. Contacts are shown using dashed lines and the contact frequency is represented by the line thickness. In the bound state, F37^CGRP^ plays a key anchoring role between CLR and RAMP by forming hydrophobic interactions with W84^RAMP1^, P85^RAMP1^, F83^RAMP1^ and W72^CLR^ and Y124^CLR^. The amide group of F37^CGRP^ also creates a strong hydrogen bond with the main T122^CLR^ main chain and a less frequent hydrogen bond with the main chain of W121^CLR^. A36^CGRP^ engages with the W121^CLR^ residue and, in addition, R119 located in the CLR turret loop via backbone hydrogen bonding. K35^CGRP^ also engages R119 through backbone hydrogen bonding, while S34^CGRP^ forms a hydrogen bond with another residue in the CLR turret loop, H114. G33^CGRP^ creates a very persistent backbone hydrogen bond with W121^CLR^ and, at the following patch, V32^CGRP^ generates extensive contacts with four CLR residues, Y124, T125, N128, F95. T30 serves as a central node of the peptide by forming hydrophobic contacts with the CLR residues D94 and F95 on *β* 3 − *β* 4 loop, and W72 - the Trp shelf. F27^CGRP^ also engages the *β* 3 − *β* 4 loop residue D94 with low frequency.

We identified a set of high-probability interactions common between both peptides: G33^CGRP^–W121^CLR^–T122^CLR^–F37^CGRP^– Y124^CLR^–V32^CGRP^–N/D31^CGRP^–T30^CGRP^–D94^CLR^-F95^CLR^. We refer to this set as the “central cascade”. This core set of stabilizing interactions solidifies the peptide conformation and bridges the CLR turret loop, the CLR *β* 3 − *β* 4 loop, and the *α*2-*α*3 loop of RAMP1. The central cascade is also preserved in the bound state of ssCGRP(27-37), with frequencies comparable to those of CGRP(27-37). The key differences between the peptides emerge in the tail region, where the substituted residues are located. While S34^CGRP^ interacts with H114^CLR^ with moderate frequency, P34^ssCGRP^ interacts with the same residue less frequently, instead strongly interacting with S117^CLR^, a contact absent in the CGRP-bound state. This interaction appears to initiate a new anchoring path as S117^CLR^ further interacts with W35^ssCGRP^, which, in addition to F37, serves as a second RAMP1 anchoring residue by interacting with F83^RAMP1^.

In the near-bound state of CGRP(27-37), the core interaction cascade at the C-terminus remains largely intact relative to the bound state, up to V32^CGRP^. V32^CGRP^ forms a new stable contact with W72^CLR^, while T30^CGRP^ loses its interaction with the *β* 3 − *β* 4 loop residues of CLR. Instead, the N-terminal segment reorganizes to form new interactions with V28^CGRP^ and F27^CGRP^ interacting with the CLR *β* 3 − *β* 4 loop residues F92 and D94, respectively. Within the tail region, a notable shift is observed where the interaction between A36^CGRP^ and R119^CLR^ becomes stronger, whereas F37^CGRP^ completely loses its already weak contact with W121^CLR^. In contrast, the near-bound state of ssCGRP(27–37) retains the central interaction cascade and even features a slightly expanded contact network. Importantly, ssCGRP simultaneously engages all key turret loop residues—H114, S117, N118, and R119—a feature not achieved by CGRP. The inability of K35^CGRP^ to interact with N118^CLR^ is likely due to electrostatic repulsion between the two positively charged side chains of R119^CLR^ and K35^CGRP^. This generally enhanced network of C-terminal interactions for ssCGRP may explain the enhanced interaction between W35^ssCGRP^ and F83^RAMP1^, a key anchoring point at the CLR–RAMP1 interface.

In addition to stable intermolecular contacts with the receptor, the intrinsic conformational stability of the peptide itself plays a critical role in its dissociation behavior. By analyzing intrapeptide contact frequencies (Supplementary Fig. 5), we observed that ssCGRP exhibits enhanced internal stability along unbinding pathways compared to CGRP. Particularly in the bound and near-bound states, the sidechain hydroxyl group of S36^ssCGRP^ generates a strong hydrogen bond with the backbone oxygen of D31^ssCGRP^, which in CGRP is replaced with a weaker interaction between backbone-backbone of A36^CGRP^–N31^CGRP^. This stabilization coincides with a general strengthening of the beta-turn motif. Strikingly, ssCGRP exhibits more intramolecular structure in the full unbound state than in the transition state. This underscores how the stabilization of internal structure is related to the prolonged residence time of ssCGRP. Stabilizing internal interactions must be broken in order to reach the ligand unbinding transition state. This adds to the already steep energetic barrier along the unbinding transition path.

## Discussion

By combining both BLI data and extensive WE-based REVO unbinding simulations, we have generated mechanistic insight into the unbinding processes for native CGRP and a high-affinity variant ssCGRP in the CLR:RAMP1 receptor complex. BLI experiments revealed that ssCGRP(8-37) has a markedly slower dissociation rate by a factor of 430, and a faster association rate by a factor of 4.5 compared to CGRP(8–37), resulting in a binding affinity ≈ 2000-fold tighter. The corresponding residence time for ssCGRP(8-37) is 132 s, in comparison with 0.3 s for CGRP(8-37). In REVO simulations of the 8–37 fragments, CGRP(8–37) yielded only a single unbinding event while ssCGRP(8–37) did not unbind, precluding reliable rate estimates. However, for the shorter 27–37 peptides, residence times of 7.5 *µ*s and 20 ms are obtained for CGRP(27-37) and ssCGRP(27-37), respectively. This corresponds to an deceleration factor of approximately 2700 for ssCGRP compared to CGRP, which is qualitatively in line with experimental results, but shows a stronger deceleration by roughly a factor of 6. Several methodological factors could explain the discrepancies between simulation and experimental dissociation rates. In BLI experiments, the N-terminal portion of the peptide in the 8–37 constructs is sterically constrained and unable to move freely. Our 8-37 simulations attempted to model this constraint using a set of repulsive potentials that prevented the N-terminus from interacting with the receptor. In the 27–37 simulations, the N-terminus is unconstrained. This difference could potentially impact peptide flexibility, thereby affecting the residence time. Our results also emphasize the importance of clearly defining what constitutes an unbound state — particularly in systems where peptides have a high potential for rebinding to the receptor after transient unbinding events. Since BLI measures dissociation by detecting complete loss of binding at the surface, simulation definitions of unbinding should aim to reflect this irreversible disengagement. Our unbound state in the WE simulations (*>* 10 Å minimum ligand-receptor atomic distance) was chosen to correspond to the lack of nonbonded interactions between the ligand and the receptor. Further investigation is warranted to determine the probability of rebinding of these states, although this would necessitate much larger simulation boxes, with increased computational cost. Finally, prior studies have shown that force field choice can influence protein–peptide binding kinetics^54^. Despite these challenges, our simulations consistently captured the prolonged residence time of ssCGRP and we are confident that valuable insights can be drawn from their modeling of the atomic interactions in the bound and near-bound states.

Our contact map analysis in the bound state reveals a set of stabilizing interactions between CGRP or ssCGRP and the CLR:RAMP1 ECD that are largely consistent with previous structural and MD studies. Notably, structural and short-timescale MD studies have highlighted persistent interactions between CGRP/ssCGRP T30 and CLR D94, and contributions of CGRP/ssCGRP V32 and T30 interacting with CLR Q93 and W72^14,15^. The *β* 3 − *β* 4 loop residues, including D90, D94, and Q93, have also been identified as important contributors to peptide association in CGRP binding simulations^55^. Our findings further explain the importance of these interactions by showing that they participate in a stable interaction cascade in the bound state of both peptides, which is largely preserved in the near-bound ensemble. This cascade also includes the stable beta-turn structure that must be disrupted for unbinding, which introduces an additional energetic barrier to dissociation that is notably higher for ssCGRP. For the C-terminal region of the CGRP peptide, interactions between CGRP F37 and CLR W121 and T122, as well as hydrogen bonds involving peptide residues G33–S36/A36 and CLR residues S117, R119, and W121, have been identified as key stabilizing contacts^14,15^. Our simulations agree with these findings and additionally reveal that ssCGRP W35 also forms a contact with CLR N118, a turret loop residue not contacted by CGRP, suggesting that complete engagement of the turret loop (H114, S117, N118, R119) is a distinguishing feature of ssCGRP. With respect to RAMP1 interactions, it has been previously noted that CGRP lacks persistent hydrogen bonds and instead engages in hydrophobic interactions, particularly involving peptide F37 and RAMP1 W84^14–16^. In addition to W84, which has been previously described as a critical residue that completes the anchoring pocket for peptide C-terminal engagement, our data also suggest an active role for RAMP1 F83. In our simulations, F83 consistently forms strong interactions with ssCGRP W35 in bound and near-bound ensembles, a pattern not observed for CGRP. While F83 has been previously noted in the context of interactions with CLR R199, our findings suggest an additional role in peptide stabilization.

Our simulations for CGRP(27–37) and ssCGRP(27–37) reveal critical distinctions in addition to specific interactions that could only be shown by extensive sampling of unbinding pathways. Detailed analysis of these pathways has shown many key differences: (I) In the bound state, ssCGRP(27–37) forms a stronger network of stabilizing interactions, especially with turret loop residues in the CLR and the critical RAMP1 F83 side chain. (II) The C-terminal substitutions of ssCGRP stabilize a beta-turn motif that must be disrupted for unbinding, creating a higher barrier to dissociation. (III) ssCGRP samples a more heterogeneous ensemble of conformations in the 27–32 region in the bound state, reflecting an entropic broadening of the energy funnel. (IV) In the near-bound state, ssCGRP retains persistent interactions, especially in the 32–37 tail region, generating a “sticky” intermediate that slows complete dissociation. (V) ssCGRP demonstrates a higher degree of ligand recapture of intermediate unbinding conformations, which fall back towards the fully bound state with high probability. Together, these differences are indicative of a more steeply funneled free energy landscape, with the bound state located in a deep, narrow well at the base and the funnel progressively widening as the ligand progresses toward the unbound ensemble. Enhanced turret loop and RAMP1 F83 contacts (I) together with more stable C-terminal beta-turn (II) deepen the free-energy funnel for ssCGRP, which makes escaping the funnel thermodynamically more costly. Similarly, this is true for a more heterogeneous ensemble of bound conformations (III), which entropically lowers the free energy of the bound state. That the stabilizing effect of the ssCGRP interaction network persists into the near-bound conformations (IV) demonstrates that this free-energy funnel is deepened not only in the middle, but more broadly. A steeper funnel could also lead to more frequent ligand recapture at intermediate conformations (V), which are halfway up the funnel wall. Together, these observations provide a refined view of peptide recognition in the CLR:RAMP1 complex.

To the best of our knowledge, this is the first study to perform a comparative mechanistic analysis of peptide ligand dissociation from a protein target. This absence is especially striking given the pharmacological importance of peptide therapeutics. Peptides offer unique advantages over antibodies or small molecules by combining biologic-level potency with advantages such as structural complexity, low toxicity, reduced immunogenicity, low production cost, and even drug-like oral availability due to recent innovations^35^. Peptides occupy a distinct niche between small molecules and biologics, offering better fit for certain targets. A comparative analysis of peptide ligand dissociation can build a working understanding of interactions throughout the unbinding pathway, providing insight into the relationship between the structural complexity and residence time. Identifying opportunities for further improvement of peptide therapeutics by providing mechanistic insight can inform rational drug design strategies not only for peptides but also for small-molecule and antibody therapeutics. The small molecule antagonists olcegepant and telcagepant inhibit CGRP binding by occupying the F37 binding site and forming hydrogen bonds with CLR T122. The olcegepant antagonist additionally binds CLR D94, F92 and W72, and the binding of both molecules shift RAMP1 *α*2 closer to CLR, stabilized by interactions with W74 and W84^32^. These interactions are generally consistent with small shifts that occur upon binding of other FDA-approved next-generation gepants targeting the CLR:RAMP1 complex^33,56^. The monoclonal antibody erenumab binds to several already mentioned key CLR residues including R119, S117, T122, W72, and D94, and RAMP1 residues W84, W74, and D71^31^. Notably, disruption of hydrogen bonds with S117 and R119 resulted in a fivefold increase in dissociation rate of erenumab. Although prior structural and MD studies have highlighted many of the same residues, the dynamic nature of peptide ligands means that static structures often miss the persistence and temporal coordination of these interactions during unbinding. Our findings reinforce that an effective strategy for designing long-acting antagonists should aim to preserve the critical interaction cascade that we have identified. These contacts are not only consistent with key interactions observed in approved therapeutics involving CLR residues D94, W72, and T122, but also extended by our dynamic analysis, which reveal their role in stabilizing bound and near-bound conformational ensembles. In particular, the 27–32 region of the peptide remains a largely under-exploited contact zone and our simulations suggest that increased interactions in this region, especially effecting the *β* 3 − *β* 4 loop, could complement central cascade stabilization to further prolong unbinding kinetics. Additionally, our simulations show that the involvement of all CLR turret loop residues could be correlated with a more stable structure and prolonged residence time. This suggests that ligands capable of stabilizing this loop through multi-point anchoring are likely to exhibit improved pharmacokinetic profiles and to date, no approved or experimentally designed CGRP-targeting therapeutics have been shown to interact with all four of these turret loop residues simultaneously, highlighting a promising and underexplored design strategy. Moreover, RAMP1 F83 emerges as a critical discriminator of ssCGRP binding, forming strong and persistent interactions with W35 that are absent in native CGRP and provide an additional anchor at the RAMP1 interface.

Several aspects of this work are broadly generalizable to other systems, particularly in contexts where dynamic mechanisms govern functional outcomes. Crucially, capturing these features would not have been possible without the use of enhanced sampling techniques such as weighted-ensemble (WE) simulations with the REVO algorithm, which enabled us to efficiently generate unbinding trajectories that occur on long timescales. This method was essential to discover the critical mechanistic features of ligand (un)binding that contribute to a prolonged residence time. While this approach may not be suitable for every system, it is particularly well-suited to those characterized by slow dissociation kinetics, entropically broadened energy funnels, or complex conformational landscapes. As these characteristics are common for many ligand–receptor interactions, the broader adoption of REVO and similar path-sampling methods could support the rational design of long residence time peptide therapeutics. Second, the design principles that emerge from our analysis can guide peptide optimization for other GPCRs and beyond. The stabilization of a beta-turn at the C-terminus emerged as a particularly potent mechanism of kinetic enhancement, acting as a structural energy barrier that slows dissociation. General strategies that constrain peptide flexibility, such as the incorporation of stabilizing secondary structural motifs like beta-turns or cyclization, can reduce conformational entropy and promote longer residence times^57^. In our study, the overall stabilized structure not only hindered complete unbinding but also supported frequent ligand recapture events, where the peptide re-engages the binding site after partial dissociation. This suggests that despite their intrinsic flexibility, peptides can maintain persistent and functionally critical interactions when specific stabilizing features are embedded in their design. Taken together, these results provide insight that can guide rational peptide design to achieve improved kinetic and pharmacological profiles. Next-generation peptide therapeutics can be tuned for kinetic selectivity by having a prolonged residence time, a higher probability of ligand recapture, and specific engagement of transient intermediate states.

## Methods

### System preparation for molecular dynamics simulations

In this study, four separate systems are constructed. Each system includes the heterodimer complex of the ECDs of CLR and RAMP1 (CLR:RAMP1) and one of four peptide constructs: CGRP(27-37), ssCGRP(27-37), CGRP(8-37), ssCGRP(8-37). The 27-37 constructs only contain the ECD-binding segment of the peptide, while the 8-37 constructs also contain part of the TMD-binding segment of the peptide as well. A comparison of the 8-37 and 27-37 peptide structures is shown in Supplementary Fig. 3. Note that CGRP has two forms in humans, *α*-CGRP and *β*-CGRP, and in this study, CGRP always refers to the *α*-CGRP isoform, the predominant form in the central and peripheral nervous systems.

For systems in which only the (27-37) segment of the ligand is included, the crystal structure of the ssCGRP (residues 27–37) bound to CLR (residues 33–130) and RAMP1 (residues 24–111) ECDs (PDB ID: 4RWG^15^) is used as a template to generate the starting structures. The maltose binding protein included in the original crystal structure as a crystallization module (residues 3–374) is omitted in our simulations, retaining only the relevant receptor and ligand components. For CLR:RAMP1:ssCGRP(27-37), this initial receptor-ligand complex structure is used as is and for the CLR:RAMP1:CGRP(27-37),four residue mutations, D31N, P34S, W35K and S36A, are reversed to their wild-type form using the CHARMM-GUI web server^58^. For the 8-37 systems, the AlphaFold3 web server^59^ is used to generate the peptide starting structures. The CLR (residues 29–134), RAMP1 (residues 24–111), and ligand (residues 8–37) FASTA sequences are taken from the UniProt database^60^ with UniProt ID Q16602, O60894, and P06881, respectively. Mutations specific to the ssCGRP variant are introduced directly into these sequences. The structures obtained using AlphaFold3 are compared with the crystal structures PDB ID: 4RWG^15^ and PDB ID: 6E3Y^14^ to ensure the precision of the models. We ensure that the key interactions between the receptor-ligand and bound conformations are conserved across all initial systems.

For all systems, the CHARMM-GUI web server^58^ is used to introduce three glycosylation sites at the CLR residues 66, 118 and 123, each with N-linked glycosylation with a beta-D-N-acetylglucosamine (bDGlcNAc) glycan attached and to generate all systems together with scripts for minimization and equilibrium. Additionally, the three known disulfide bonds between residues C88-C127, C48-C74, and C65-C105 in CLR and residues C57-C104, C40-C72, and C27-C82 in RAMP1 are preserved in all structures. The C-termini of the peptides are amidated. Each system is solvated in the TIP3P water model that has a cutoff of up to a 10 Å from the protein to each edge of the periodic box. The length of the simulation box is set to ∼75 Å and ∼77 Å in each dimension for the 27-37 and 8-37 systems, respectively. Sodium and potassium atoms are then added to neutralize each system and achieve an ionic concentration of 150 mM. The solvated systems are ∼35,928 and ∼42,740 atoms in size for the 27-37 and 8-37 systems, respectively.

### Molecular dynamics simulations

For each system, all-atom MD simulations are performed in an explicit solvent using the OpenMM simulation engine^61^ along with the CHARMM36 force field^62^. Prior to the production run, energy minimization of the systems is performed with a maximum number of 5000 steps and a minimization energy tolerance of 100 kcal/mol to remove steric clashes and stabilize the structure. The equilibration run is performed using a time step of 0.001 ps for 125000 steps with positional restraints on the protein backbone (400 kJ/mol/nm^2^) and side chains (40 kJ/mol/nm^2^), while dihedral restraints (4 kJ/mol/rad^2^) are applied to the carbohydrate moieties to minimize excessive fluctuations and preserve conformational stability. All covalent bonds with hydrogens are also constrained. Subsequently, the protein restraints are removed and the production run is performed using a 0.002 ps time step, with a total simulation length of 10 ns. Both equilibrium and production runs are performed under constant particle number (N), constant temperature (T) and constant pressure (P) ensemble conditions, where the temperature was maintained at 310 K using Langevin dynamics with a friction coefficient of 1 ps^−1^ to ensure efficient temperature control and constant pressure is maintained at 1.0 bar using an isotropic Monte Carlo barostat with coupling frequency being performed every 100 steps. Long-range electrostatic interactions are treated using the particle mesh Ewald method, with an Ewald error tolerance of 0.0005. The van der Waals interactions are calculated using the force-switch method, with a switching distance of 10 Å and a cutoff distance of 12 Å. Periodic boundary conditions are applied.

For the 8-37 systems, a custom centroid bond force is implemented at the N-terminus of the peptides during production runs to ensure that the TMD-binding segment of the ligands would not directly interact with the receptor complex. This mimics the experimental conditions as this part of the peptide is further extended by three additional residues and biotinylated in the BLI assays (see the experimental determination of rates section in Methods). The centroid bond force takes the form of a purely repulsive potential that is activated only when the distance between groups (*r*) is smaller than a cutoff (*r*_0_). We use a number of groups on CLR and RAMP1, including K134, P97, and P90 in the CLR structure, the center of mass of the N-terminal helix of CLR, center of mass of CLR, center of mass of the N-terminus of RAMP1, and the center of mass of RAMP1. For each of these groups, we use a separate *r*_0_ value that was determined using the center of mass distances in the initial structure (Table 2). We also include a direct constraint that was enforced between residues R11 in ligands and S98 in CLR due to their close proximity. The form of the repulsive potential is as follows:

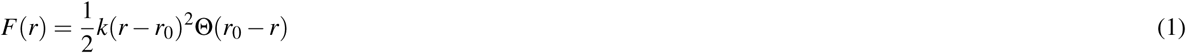

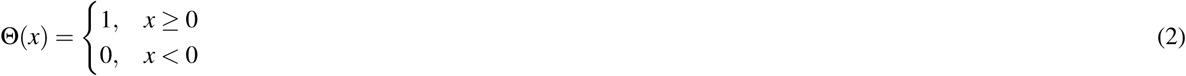

where *k* is the force constant defining the strength of the restraint and set to 20 kJ/mol/nm^2^, *r*_0_ is the cutoff distance between the selected groups and Θ(*x*) ensures that the restraint only applies when the distance falls below the threshold. The restraints respect periodic boundary conditions to maintain consistency in the simulation box.

**Table 2:** Distance cutoff values (*r*_0_) for CLR and RAMP groups. All *r*_0_ values are reported in nm. ^1^For the N-terminal helix of CLR, residues 29 to 35 were used. ^2^For the N-terminal helix of RAMP1, residues 110 and 111 were used.

| Selection | $r_0$ (CGRP) | $r_0$ (ssCGRP) | peptide selection |
| --- | --- | --- | --- |
| K134 (CLR) | 1.05 | 1.04 | 8-15 |
| P97 (CLR) | 1.30 | 1.37 | 8-15 |
| P90 (CLR) | 1.50 | 1.58 | 8-15 |
| N-term. helix (CLR)<sup>1</sup> | 2.20 | 2.20 | 8-15 |
| Entire CLR | 2.60 | 2.68 | 8-15 |
| N-term. helix (RAMP1)<sup>2</sup> | 3.80 | 3.75 | 8-15 |
| Entire RAMP1 | 1.99 | 2.05 | 8-15 |
| S98 (CLR) | 1.19 | 1.23 | R11 |

### Ligand unbinding simulations with REVO

The final structures obtained after MD simulations are used to initialize Weighted Ensemble (WE) enhanced sampling simulations^43^. In our WE simulations, an ensemble of trajectories is initiated with equal statistical weights. As the simulation progresses, each trajectory evolves independently, exploring different conformational states. In order to increase the possibility of observing events of interest, some of these trajectories (also called “walkers”) are cloned into multiple copies. Conversely, two walkers can also be merged to conserve computational resources. Merged walkers are typically trapped in local or global free energy minima or are redundant in the ensemble. To preserve overall probability, the assigned weights are evenly divided among new clones for cloned walkers, and the weights are summed for merged walkers at each resampling step. As is true in all WE methods, a random selection process governs the merged trajectories, where the surviving walker is chosen with a probability that is proportional to its weight. The precise nature of this selection is critical to ensure that the net probability flux between regions of conformational space due to merging remains zero^63^.

The resampling of ensembles by variation optimization (REVO) algorithm^46^ is an implementation of the WE method designed for efficient and adaptive resampling by performing cloning and merging steps based on the trajectory variation (*V*). Notably, rather than relying on spatial bins, REVO identifies underexplored regions dynamically and redistributes computational resources accordingly. It calculates *V* as follows:

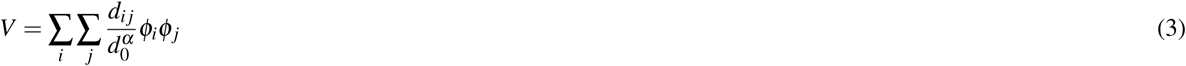

where *d_i j_* represents the distance between walkers *i* and *j*, *d*_0_ is a characteristic distance to normalize the variation, *α* is a scaling exponent (here set to 4), *φ_i_* and *φ_j_* are weighting functions for the walkers that here are solely a function of the walker weights:

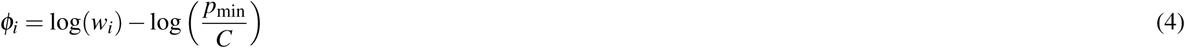

where *w_i_*is the weight of the trajectory *i*, *p*_min_ is the predefined minimum trajectory weight (set to 10^−12^), and *C* is a normalization constant set to 100, following previous studies. REVO is used as implemented in wepy^64^, which is a Python implementation of the weighted ensemble method. The resampling algorithm used in this study is guided by the `UnbindingDistance` method from wepy, which quantifies distances between walkers in the ensemble by calculating the root-mean-square deviation (RMSD) of the ligand atoms after aligning the receptor atoms to a common reference structure. The distances are then used to calculate the initial *V* to select a candidate walker with the highest individual variation and with a weight exceeding *p*_min_.

Two candidate walkers are also chosen for merging using the “pairs” algorithm in REVO, in which the pair of walkers with the lowest expected variation loss is selected that meets the following criteria: 1) the sum of their weights is less than *p*_max_, 2) their distance is below a specified merging threshold, and 3) neither of which is the cloned walker. If all three walkers are successfully identified, cloning and merging operations are proposed. If a proposed merging and cloning event results in an overall increase in *V*, it is accepted and another resampling event is proposed. This iterative process continues until *V* reaches a local maximum, at which point the resampling step concludes and the ensemble is propagated forward again in time using MD integration. To enhance the likelihood of observing unbinding events, two distinct `UnbindingDistance` selection criteria are utilized in this study. In one group of simulations, the tail region of the ligand (residues 32-37) is selected for RMSD calculation, while in the other group, the entire ligand.

Simulations are conducted in the unbinding ensemble, a non-equilibrium framework where trajectories that reach the unbound state are reset to the bound state. This is implemented using the `UnbindingBC` boundary condition method from wepy, with the unbound state is defined as when the minimum distance between the ligand and the receptor exceeds 10 Å. For the minimum distance calculation, all atoms in the ligand are used, and the receptor reference set is defined as atoms within 10 Å of the ligand in the equilibrated bound pose. The unbinding rates can be calculated directly from unbinding simulations by evaluating the flux into the unbound state. This flux is calculated using Hill’s equation^65,66^, where the sum of the weights of the trajectories reaching the unbound state is divided by the elapsed time:

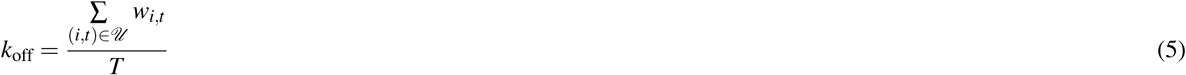

where, *U* represents the set of index-time pairs (*i, t*) corresponding to unbinding events, *w_i,t_* is the statistical weight of the trajectory *i* at time *t*, and *T* is the total elapsed time of the simulation.

Each simulation cycle consists of 20 ps of MD (10,000 steps at a time step of 0.002 ps) followed by a round of REVO resampling. 12 and 6 independent simulations were performed for each 27-37 systems and 8-37 systems, with each system having 11000 and 15250 cycles totaling 126.72 *µ*s and 87.84 *µ*s, respectively (Table 1). Each run has 48 walkers, and the chosen ensemble size of 48 walkers was selected to balance computational efficiency with sufficient trajectory diversity, ensuring broad coverage of the ligand unbinding pathways while allowing extended simulation timescales. Additionally, for efficient parallelization, the ensemble size was set to be divisible by the number of GPUs on a given node (in this case, 8). Snapshots of each trajectory were stored after each cycle to allow for detailed kinetic and structural analysis.

### Markov state modeling

Markov State Models (MSMs) are built by projecting each frame onto a set of structural features and discretizing this space into states to determine metastable and transition states. The features are a set of interatomic distances, comprised of 198 distances calculated between 18 C*α* atoms at the CLR binding site (residues G71, W72, F92, Q93, D94, F95, D114, A116, S117, R119, W121, T122, N123, Y124, T125, N128, V129, N130) and 11 C*α* atoms from the ECD-binding region of ligands, as well as 88 distances between 8 C*α* atoms of RAMP1 (residues D71, W74, E78, G81, K82, F83, W84, P85) and same 11 ligand C*α* atoms from the ECD-binding region of ligands (Supplementary Fig.4). Dimensionality reduction is performed using Time-Structure Independent Components Analysis (tICA) implemented in the Deeptime library^67^ to improve clustering efficiency and focus on kinetically relevant motions. Approximately 3 million data points are generated via the `sliding_windows` function from wepy, from which 750,000 randomly selected points are used to train the tICA model, which is then used to transform the entire set. The reduced-dimensional data are clustered into discrete states using KMeans clustering algorithm from scikit-learn^68^, assigning each trajectory frame to a cluster index. A set of MSMs are systematically built by varying the number of clusters from 50 to 2000, and the output dimensions of tICA from 3 to 10. Transition count matrices were constructed using time-lagged state assignments at a 20 ps lag time.

These matrices are utilized to construct conformation space networks (CSNs) using the CSNAnalysis package^69^. The clusters are labeled bound if the average ligand RMSD relative to the initial reference structure is less than 3.0 Å and unbound if any frame within the cluster exhibited a minimum distance greater than 8.0 Å between any atom on the ligand and CLR. There, a more relaxed definition of unbound is used than the minimum distance of 10 Å during the run time due to the flexible nature of peptide ligands. After defining bound and unbound basins, committor probabilities and mean first-passage times (MFPTs) are calculated using the CSNAnalysis functions `calc_committors` and `calc_mfpt`, respectively. MFPT calculations are performed by creating first-passage time distributions at intervals of 2*iτ* (with *τ* being the MSM lag time), continuing until 99.9% of trajectories reached the unbound basin. The unbinding committor probability, derived from the transition path theory^70^, quantifies the likelihood that an intermediate state will transition (commit) to the unbound basin before the bound basin. The committor probabilities for states outside the bound and unbound basins are in the range (0,1), where 0 being for states that are very close to the bound basin and where 1 being for states that are very close to the unbound basin.

### Experimental determination of rates

#### Cell culture

HEK293T (Human embryonic kidney cell line, female; CRL 3216) cells were from American Type Culture Collection (Manassas, VA, USA). Cells were cultured in Dulbecco’s modified Eagle’s medium (DMEM with 4.5 g/L glucose and L-glutamine and 110 mg/L sodium pyruvate) from Gibco (11995-081) with 10 % v/v fetal bovine serum (Gibco 16000-044). Cells were grown at 37°C, 5% CO2 in a humidified incubator and passaged twice per week.

#### Plasmid constructs

pHLsec/R1.24-111-(GS)5-CLR.29-134-H6 was constructed using traditional PCR and restriction enzyme cloning methods. The insert was amplified from pAP462 (R1.24-111-(GS)5-CLR.29-144-Avitag-H6) using forward primer 5’- GCT-GAAACCGGTACGACTGC CTGCCAGGAGGC-3’ (pri742) and reverse primer 5’-GTGCTTGGTACCTTAATGATGGTGGTGATGATGTTTCTCGTGGGTGTTAAC-3’ (pri650) and inserted between AgeI and KpnI sites in the pHLsec vector.

#### Synthetic peptides

Synthetic custom peptides were synthesized and HPLC-purified by RS Synthesis (Louisville, KY) are as follows: biotin-CGRP(8-37)NH2 and Biotin-CGRP(8-37)NH2[N31D/S34P/K35W /A36S]. The lyophilized powders were resuspended at 10 mg/mL in sterile ultrapure water. Concentrations of the peptides were determined by UV absorbance at 280 nm with dilutions in 10 mM Tris-HCl, 1 mM EDTA at pH 8.0. Extinction coefficients were calculated from Tyr, Trp, and cystine content. The concentration of biotin-CGRP(8-37)NH2 was determined by the peptide content reported from RS Synthesis. Peptides were stored at −80°C with small aliquots to limit the number of freeze-thaws.

#### Protein expression and purification

N-glycosylated fusion protein was expressed similar to described [Roehrkasse, 2018, JBC] in HEK293T cells. The cells were grown in 8 or 10 T175 flasks to 80-90% confluency in DMEM supplemented with 10% fetal bovine serum at 37°C and 5% CO2. Cells were transiently transfected with 50 µg plasmid DNA and 75 µg polyethylenemine (PEI) per flask. RAMP1.24-111-(GS)5-CLR.29-134-H6 was expressed for 3 days at 37°C and 5% CO2. Culture media was harvested and filtered with 0.45 µm filter before the proteins were purified by immobilized nickel affinity chromatography followed by size exclusion chromatography. The protein was stored on ice in 50 mM Tris-HCl pH 7.5 and 150 mM NaCl until BLI could be performed the following day. The purified protein was quantitated using UV spectroscopy with a yield of 7 mg for 8 T175 prep and 10 mg for 10 T175 prep.

#### Biolayer Interferometry (BLI)

Binding kinetic experiments using the RAMP1-CLR ECD purified fusion protein were done on the Octet RED96 instrument using high precision streptavidin (SAX) biosensors from Sartorius (Göttingen, Germany). The biosensors were equilibrated in 200 µL of the assay buffer overnight at 4°C. All experiments were performed at 25°C and 1,000 rpm. Experiments with biotin-CGRP(8-37)NH2 were conducted in 50 mM Tris-HCl at pH 7.5, 150 mM NaCl, 5 mM KCl, 2 mM CaCl2, 5 mg/mL fatty-acid-free bovine serum albumin (FAF-BSA), and 0.05% tween-20 at 10 Hz. A baseline of the biosensors was taken in assay buffer for 5 minutes followed by immobilization of the biotinylated peptide at 20 nM for 200 seconds and then another baseline in buffer for 2 minutes. The biosensors were then dipped in 2-fold serial dilutions of purified RAMP1-CLR ECD fusion protein starting at 50 µM for 1 minute. Dissociation was initiated by dipping the biosensors back into buffer for 1 min. Experiments with biotin-CGRP(8-37)NH2 [N31D/S34P/K35W/A36S] were conducted in 25 mM NaHEPES at pH 7.5, 150 mM NaCl, 5 mM KCl, 2 mM CaCl2, 1 mg/mL FAF-BSA, and 0.05% tween-20 at 5 Hz. A baseline of biosensors was taken in assay buffer for 5 minutes followed by immobilization of the biotinylated peptide at 5 nM for 200 seconds and then another baseline in buffer for 2 minutes. The biosensors were then dipped in 2-fold serial dilutions of purified RAMP1-CLR ECD fusion protein starting at 200 nM for 200 seconds. Dissociation was initiated by dipping the biosensors back into buffer for 10 min. Data was analyzed using the ForteBio Data Analysis software v. 9.0 with global association and dissociation fit with linked biosensors. The data was also analyzed in GraphPad Prism (v. 10.0) to make the figures. The baseline after immobilization was subtracted followed by another baseline subtraction of the control biosensor. The time was adjusted t=0 for the association phase. The curves were then fit to the association then dissociation equation in GraphPad Prism. GraphPad Prism was only used to make the figures presented, and ForteBio Data Analysis software was used for the reported values.

## Acknowledgments

This work was supported by National Institutes of Health grants R01GM104251 (A.P.) and R01GM130794 (A.D.). Use of the Octet Red biolayer interferometry instrument at the OU Protein Production and Characterization Core facility was supported by Institutional Development Awards (IDeA) from the National Institute of General Medical Sciences of the National Institutes of Health (Grants P20GM103640 and P30GM145423), the OU Vice President for Research and Partnerships, and the OU College of Arts and Sciences.

## Author contributions

A.D. and A.P. designed the project; C.K. and A.D. collected and analyzed the computational data; K.B and A.P. collected and analyzed the experimental data; all authors prepared and reviewed the final manuscript.

## Data and code availability

The pipeline to run REVO simulations in wepy and build MSMs from weighted-ensemble data can be found in the WepyAnalysis repository at https://github.com/ADicksonLab/wepy-analysis.git. The REVO simulation data set can be downloaded from the Zenodo repository at https://zenodo.org/records/15361245. All relevant data are available from the authors upon request.

## References

1. Davis, R. B., Kechele, D. O., Blakeney, E. S., Pawlak, J. B. & Caron, K. M. Lymphatic deletion of calcitonin receptor–like receptor exacerbates intestinal inflammation. JCI insight 2, e92465 (2017).

2. Brain, S. D. & Grant, A. D. Vascular actions of calcitonin gene-related peptide and adrenomedullin. Physiol. reviews 84, 903–934 (2004).

3. Iyengar, S., Ossipov, M. H. & Johnson, K. W. The role of calcitonin gene–related peptide in peripheral and central pain mechanisms including migraine. Pain 158, 543–559 (2017).

4. Hay, D. L., Garelja, M. L., Poyner, D. R. & Walker, C. S. Update on the pharmacology of calcitonin/cgrp family of peptides: Iuphar review 25. Br. journal pharmacology 175, 3–17 (2018).

5. Russo, A. F. Calcitonin gene-related peptide (cgrp): a new target for migraine. Annu. review pharmacology toxicology 55, 533–552 (2015).

6. Argunhan, F. & Brain, S. D. The vascular-dependent and-independent actions of calcitonin gene-related peptide in cardiovascular disease. Front. Physiol. 13, 833645 (2022).

7. Udit, S., Blake, K. & Chiu, I. M. Somatosensory and autonomic neuronal regulation of the immune response. Nat. Rev. Neurosci. 23, 157–171 (2022).

8. Angenendt, L. et al. The neuropeptide receptor calcitonin receptor-like (calcrl) is a potential therapeutic target in acute myeloid leukemia. Leukemia 33, 2830–2841 (2019).

9. Kim, Y. J. & Granstein, R. D. Roles of calcitonin gene-related peptide in the skin, and other physiological and pathophysiological functions. Brain, Behav. & Immunity-Health 18, 100361 (2021).

10. McLatchie, L. M. et al. Ramps regulate the transport and ligand specificity of the calcitonin-receptor-like receptor. Nature 393, 333–339 (1998).

11. Hay, D. L. & Pioszak, A. A. Receptor activity-modifying proteins (ramps): new insights and roles. Annu. review pharmacology toxicology 56, 469–487 (2016).

12. Pal, K., Melcher, K. & Xu, H. E. Structure and mechanism for recognition of peptide hormones by class b g-protein-coupled receptors. Acta pharmacologica sinica 33, 300–311 (2012).

13. Wootten, D., Christopoulos, A., Marti-Solano, M., Babu, M. M. & Sexton, P. M. Mechanisms of signalling and biased agonism in g protein-coupled receptors. Nat. reviews Mol. cell biology 19, 638–653 (2018).

14. Liang, Y.-L. et al. Cryo-em structure of the active, gs-protein complexed, human cgrp receptor. Nature 561, 492–497 (2018).

15. Booe, J. M. et al. Structural basis for receptor activity-modifying protein-dependent selective peptide recognition by a g protein-coupled receptor. Mol. cell 58, 1040–1052 (2015).

16. Josephs, T. M. et al. Structure and dynamics of the cgrp receptor in apo and peptide-bound forms. Science 372, eabf7258 (2021).

17. Maton, P., Pradhan, T., Zhou, Z.-C., Gardner, J. & Jensen, R. Activities of calcitonin gene-related peptide (cgrp) and related peptides at the cgrp receptor. Peptides 11, 485–489 (1990).

18. Luo, G. et al. Discovery of (5 s, 6 s, 9 r)-5-amino-6-(2, 3-difluorophenyl)-6, 7, 8, 9-tetrahydro-5 h-cyclohepta [b] pyridin-9-yl 4-(2-oxo-2, 3-dihydro-1 h-imidazo [4, 5-b] pyridin-1-yl) piperidine-1-carboxylate (bms-927711): An oral calcitonin gene-related peptide (cgrp) antagonist in clinical trials for treating migraine. J. medicinal chemistry 55, 10644–10651 (2012).

19. Blair, H. A. Rimegepant: a review in the acute treatment and preventive treatment of migraine. CNS drugs 37, 255–265 (2023).

20. Voss, T. et al. A phase iib randomized, double-blind, placebo-controlled trial of ubrogepant for the acute treatment of migraine. Cephalalgia 36, 887–898 (2016).

21. Scott, L. J. Ubrogepant: first approval. Drugs 80, 323–328 (2020).

22. Deeks, E. D. Atogepant: first approval. Drugs 1–6 (2022).

23. Scuteri, D., Tarsitano, A., Tonin, P., Bagetta, G. & Corasaniti, M. T. Focus on zavegepant: the first intranasal third-generation gepant. Pain management 12, 879–885 (2022).

24. Shi, L. et al. Pharmacologic characterization of amg 334, a potent and selective human monoclonal antibody against the calcitonin gene-related peptide receptor. The J. pharmacology experimental therapeutics 356, 223–231 (2016).

25. Stoker, K. & Baker, D. E. Erenumab-aooe. Hosp. Pharm. 53, 363–368 (2018).

26. Dhillon, S. Eptinezumab: first approval. Drugs 80, 733–739 (2020).

27. Benschop, R. et al. Development of a novel antibody to calcitonin gene-related peptide for the treatment of osteoarthritis-related pain. Osteoarthr. Cartil. 22, 578–585 (2014).

28. Lamb, Y. N. Galcanezumab: first global approval. Drugs 78, 1769–1775 (2018).

29. Bigal, M. E. et al. Tev-48125 for the preventive treatment of chronic migraine: efficacy at early time points. Neurology 87, 41–48 (2016).

30. Bigal, M. E. et al. From lbr-101 to fremanezumab for migraine. CNS drugs 32, 1025–1037 (2018).

31. Garces, F. et al. Molecular insight into recognition of the cgrpr complex by migraine prevention therapy aimovig (erenumab). Cell Reports 30, 1714–1723 (2020).

32. ter Haar, E., et al. Crystal structure of the ectodomain complex of the cgrp receptor, a class-b gpcr, reveals the site of drug antagonism. Structure 18, 1083–1093 (2010).

33. Leung, L., Liao, S. & Wu, C. To probe the binding interactions between two fda approved migraine drugs (ubrogepant and rimegepant) and calcitonin-gene related peptide receptor (cgrpr) using molecular dynamics simulations. ACS chemical neuroscience 12, 2629–2642 (2021).

34. David, L., Scalley-Kim, M., Olland, A., White, A. & Misura, K. The eptinezumab: Cgrp complex structure–the role of conformational changes in binding stabilization. Bioengineered 12, 11076–11086 (2021).

35. Muttenthaler, M., King, G. F., Adams, D. J. & Alewood, P. F. Trends in peptide drug discovery. Nat. reviews Drug discovery 20, 309–325 (2021).

36. Miranda, L. P. et al. Identification of potent, selective, and metabolically stable peptide antagonists to the calcitonin gene-related peptide (cgrp) receptor. J. medicinal chemistry 51, 7889–7897 (2008).

37. Miranda, L. P. et al. Peptide antagonists of the calcitonin gene-related peptide (cgrp) receptor with improved pharmacokinetics and pharmacodynamics. Pept. Sci. 100, 422–430 (2013).

38. Srinivasan, K. et al. Pharmacological, pharmacokinetic, pharmacodynamic and physicochemical characterization of fe 205030: a potent, fast acting, injectable cgrp receptor antagonist for the treatment of acute episodic migraine. J. Pharm. Sci. 111, 247–261 (2022).

39. Jamaluddin, A. et al. Lipidated calcitonin gene-related peptide (cgrp) peptide antagonists retain cgrp receptor activity and attenuate cgrp action in vivo. Front. Pharmacol. 13, 832589 (2022).

40. Kristensen, J. B. et al. Pipeline for development of acylated peptide based cgrp receptor antagonist with extended half-life for migraine treatment. Sci. Reports 15, 1870 (2025).

41. Booe, J. M., Warner, M. L., Roehrkasse, A. M., Hay, D. L. & Pioszak, A. A. Probing the mechanism of receptor activity–modifying protein modulation of gpcr ligand selectivity through rational design of potent adrenomedullin and calcitonin gene-related peptide antagonists. Mol. pharmacology 93, 355–367 (2018).

42. Babin, K. M., Kilinc, C., Gostynska, S. E., Dickson, A. & Pioszak, A. A. Characterization of the two-domain peptide binding mechanism of the human cgrp receptor for cgrp and the ultrahigh affinity sscgrp variant. Biochemistry (2025).

43. Huber, G. A. & Kim, S. Weighted-ensemble brownian dynamics simulations for protein association reactions. Biophys. journal 70, 97–110 (1996).

44. Chong, L. T., Saglam, A. S. & Zuckerman, D. M. Path-sampling strategies for simulating rare events in biomolecular systems. Curr. opinion structural biology 43, 88–94 (2017).

45. Hénin, J., Lelièvre, T., Shirts, M. R., Valsson, O. & Delemotte, L. Enhanced sampling methods for molecular dynamics simulations. arXiv preprint arXiv:2202.04164 (2022).

46. Donyapour, N., Roussey, N. M. & Dickson, A. Revo: Resampling of ensembles by variation optimization. The J. chemical physics 150 (2019).

47. Chodera, J. D. & Noé, F. Markov state models of biomolecular conformational dynamics. Curr. opinion structural biology 25, 135–144 (2014).

48. Husic, B. E. & Pande, V. S. Markov state models: From an art to a science. J. Am. Chem. Soc. 140, 2386–2396 (2018).

49. Bose, S. et al. How robust is the ligand binding transition state? J. Am. Chem. Soc. 145, 25318–25331 (2023).

50. Dickson, A. & Lotz, S. D. Multiple ligand unbinding pathways and ligand-induced destabilization revealed by wexplore. Biophys. journal 112, 620–629 (2017).

51. Lotz, S. D. & Dickson, A. Unbiased molecular dynamics of 11 min timescale drug unbinding reveals transition state stabilizing interactions. J. Am. Chem. Soc. 140, 618–628 (2018).

52. Moad, H. E. & Pioszak, A. A. Selective cgrp and adrenomedullin peptide binding by tethered ramp-calcitonin receptor-like receptor extracellular domain fusion proteins. Protein Sci. 22, 1775–1785 (2013).

53. Roehrkasse, A. M., Booe, J. M., Lee, S.-M., Warner, M. L. & Pioszak, A. A. Structure–function analyses reveal a triple *β*-turn receptor-bound conformation of adrenomedullin 2/intermedin and enable peptide antagonist design. J. Biol. Chem. 293, 15840–15854 (2018).

54. Ahmad, K. et al. Enhanced-sampling simulations for the estimation of ligand binding kinetics: current status and perspective. Front. molecular biosciences 9, 899805 (2022).

55. Deganutti, G. et al. Exploring ligand binding to calcitonin gene-related peptide receptors. Front. Mol. Biosci. 8, 720561 (2021).

56. Schaduangrat, N., Khemawoot, P., Jiso, A., Charoenkwan, P. & Shoombuatong, W. Metacgrp is a high-precision meta-model for large-scale identification of cgrp inhibitors using multi-view information. Sci. Reports 14, 24764 (2024).

57. Bozovičar, K. & Bratkovič, T. Small and simple, yet sturdy: conformationally constrained peptides with remarkable properties. Int. journal molecular sciences 22, 1611 (2021).

58. Jo, S., Kim, T., Iyer, V. G. & Im, W. Charmm-gui: a web-based graphical user interface for charmm. J. computational chemistry 29, 1859–1865 (2008).

59. Abramson, J. et al. Accurate structure prediction of biomolecular interactions with alphafold 3. Nature 630, 493–500 (2024).

60. Uniprot: the universal protein knowledgebase in 2025. Nucleic Acids Res. 53, D609–D617 (2025).

61. Eastman, P. et al. Openmm 7: Rapid development of high performance algorithms for molecular dynamics. PLoS computational biology 13, e1005659 (2017).

62. Brooks, B. R. et al. Charmm: the biomolecular simulation program. J. computational chemistry 30, 1545–1614 (2009).

63. Zhang, B. W., Jasnow, D. & Zuckerman, D. M. The “weighted ensemble” path sampling method is statistically exact for a broad class of stochastic processes and binning procedures. The J. chemical physics 132 (2010).

64. Lotz, S. D. & Dickson, A. Wepy: a flexible software framework for simulating rare events with weighted ensemble resampling. ACS omega 5, 31608–31623 (2020).

65. Dickson, A., Warmflash, A. & Dinner, A. R. Separating forward and backward pathways in nonequilibrium umbrella sampling. The J. chemical physics 131 (2009).

66. Hill, T. L. Free energy transduction and biochemical cycle kinetics (Courier Corporation, 2005).

67. Hoffmann, M. et al. Deeptime: a python library for machine learning dynamical models from time series data. Mach. Learn. Sci. Technol. 3, 015009 (2021).

68. Pedregosa, F. et al. Scikit-learn: Machine learning in Python. J. Mach. Learn. Res. 12, 2825–2830 (2011).

69. Dickson, A. & Lotz, S. Csnanalysis, 2020. https://github.com/ADicksonLab/CSNAnalysis, Date Accessed: 04-09-2025.

70. Metzner, P., Schütte, C. & Vanden-Eijnden, E. Transition path theory for markov jump processes. Multiscale Model. & Simul. 7, 1192–1219 (2009).

