## Supplemental Info for "Determinants of Improved CGRP Peptide Binding Kinetics Revealed by Enhanced Molecular Simulations"

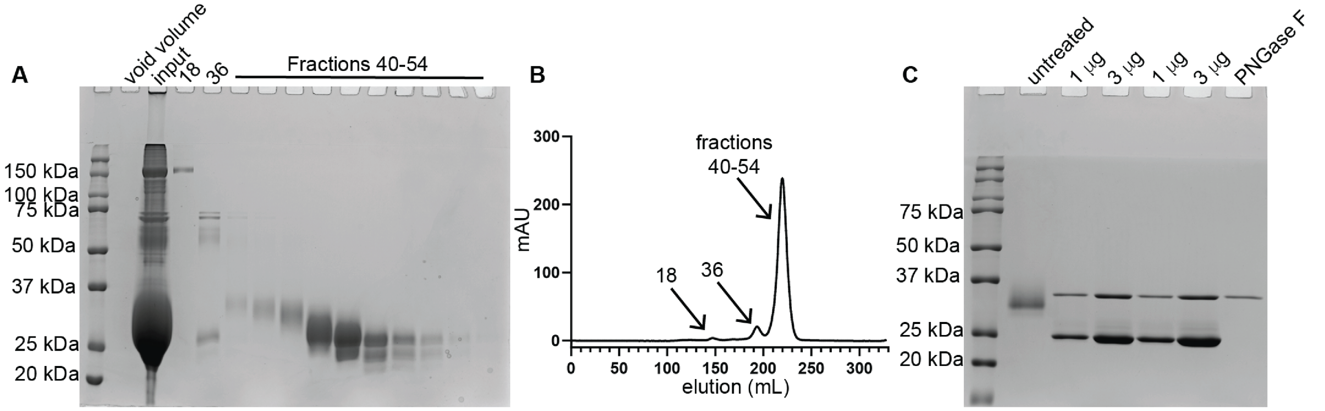


**Figure 1.** Purification and PNGase F treatment of RAMP1-CLR ECD complex. (A) 12% SDS gel with Coomassie stain of fractions collected from the size exclusion column during purification. RAMP1-CLR ECD complex is about 25 kDa. Input was the pooled fractions from the Ni column. (B) Absorbance chromatogram from the size exclusion column. (C) 12% SDS gel with Coomassie stain of purified RAMP1-CLR ECD after PNGase F treatment.

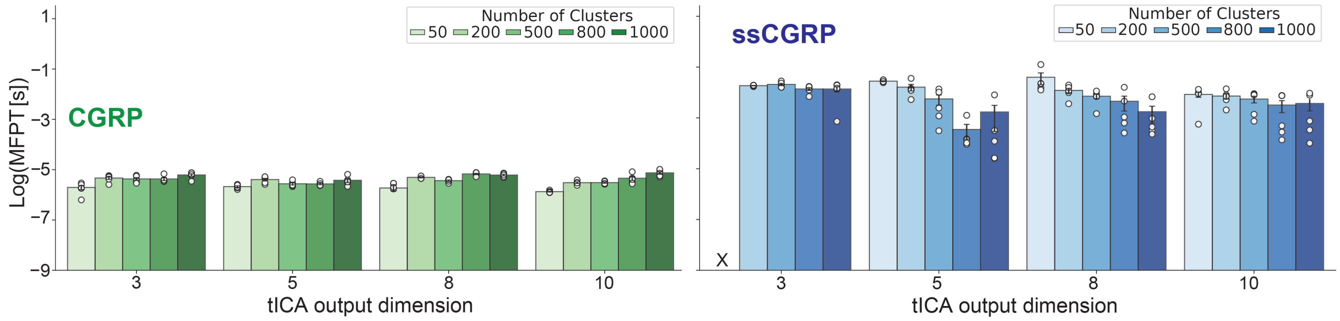


**Figure 2.** MFPT estimates with respect to the number of tICA output dimensions and number of clusters used in MSM construction. Number of tICA output dimensions ranging from 3 to 10 and number of clusters ranging from 50 to 1000 were used to calculate MFPTs (in seconds) for CGRP (left) and ssCGRP (right).

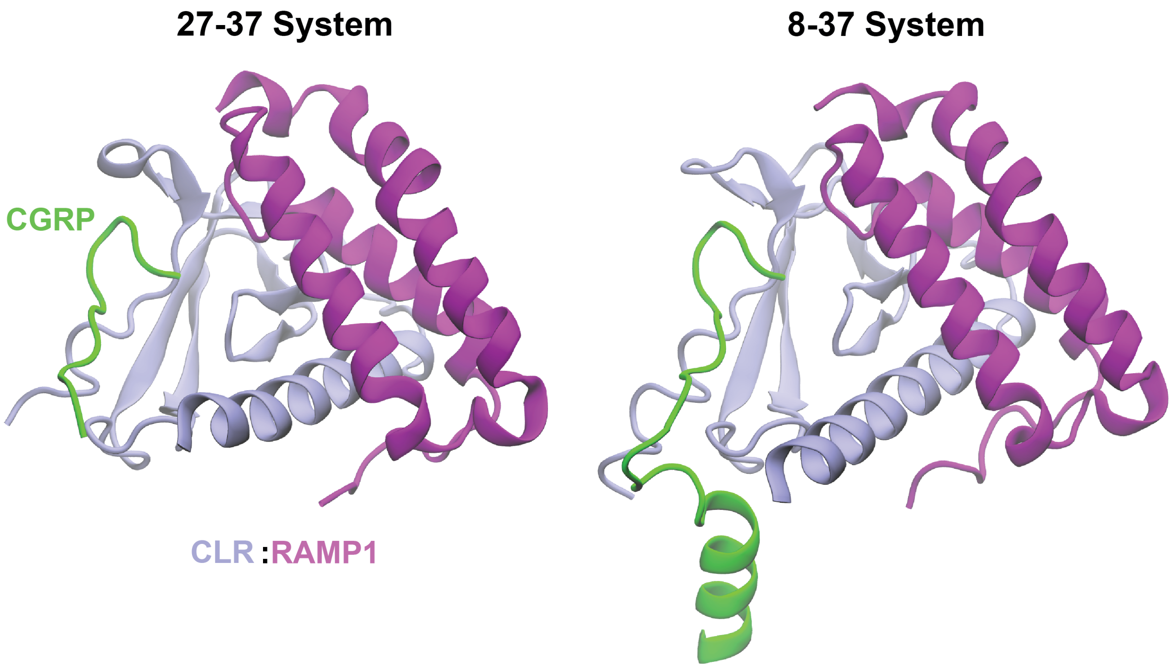


**Figure 3.** Initial structures of the (27–37) and (8–37) systems. Both the (27–37) (left) and (8–37) (right) systems include the extracellular domains (ECDs) of CLR (ice blue) and RAMP1 (purple). The (27–37) system contains only the ECD-binding segment of the peptide, while the (8–37) system additionally includes a part of the TMD-binding region of the peptide.

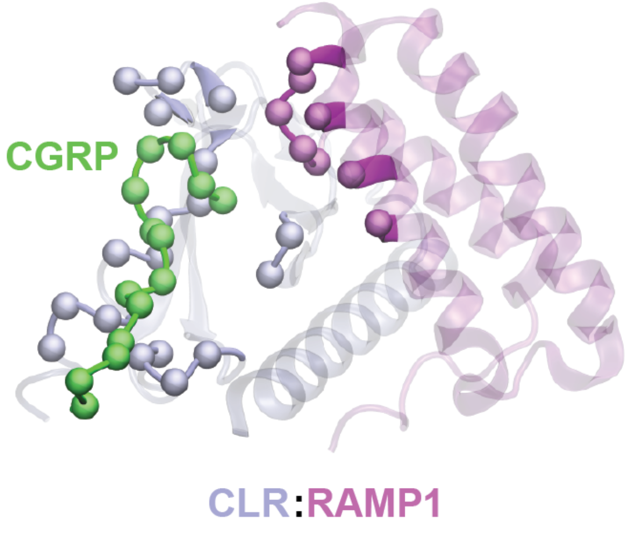


**Figure 4.** Atoms selected to calculate the distances between ligand and receptor binding site. Selected residue regions of CLR (ice blue), RAMP1 (purple) and CGRP (green) are shown in solid colors for CGRP(27-37) (left) and CGRP(27-37) (right) systems. C$\alpha$ atoms of selected regions are shown as circles.

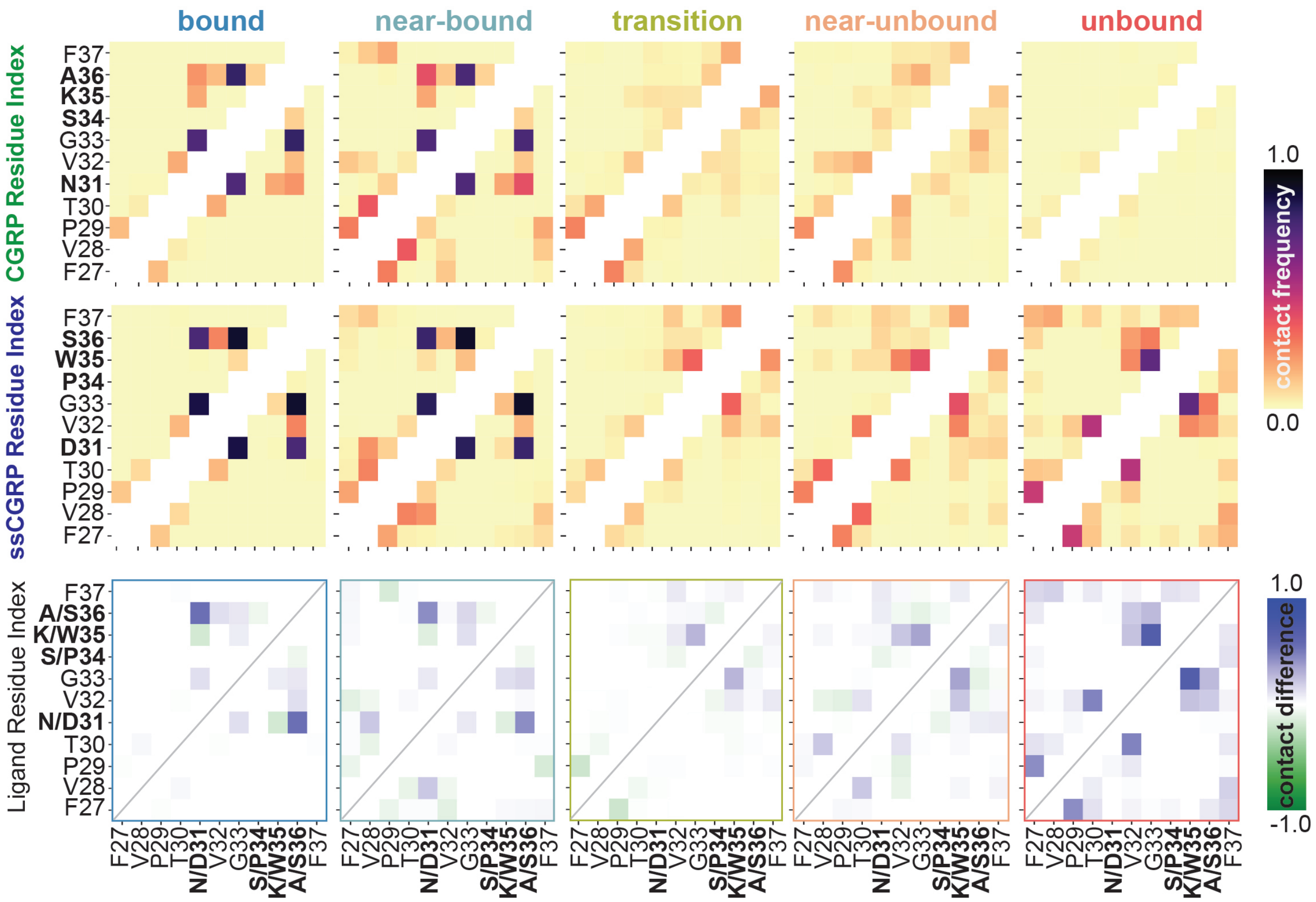


**Figure 5.** Residue-level intramolecular interaction analysis for CGRP(27–37) and ssCGRP(27–37) along their unbinding pathways. Heatmaps show the contact frequencies between peptide residues (top and middle rows) and their differences (bottom row). Warmer colors indicate more frequent contacts, while cooler tones indicate lower contact frequency. Diagonal and immediately adjacent contacts are masked in white. The contact difference map uses a diverging color scale to reflect the enrichment of specific interactions in CGRP (negative, green) or ssCGRP (positive, blue). The main diagonal is marked with a gray line for reference.

**Table 1.** Summary of kinetic values of immobilized biotin-(8–37) peptides binding purified CLR–RAMP1 ECD complexes in BLI assays. [1] Residence time calculated as the inverse of koff. [2]Half-life calculated as ln 2 divided by koff. [3] Values derived from Fig. IV-3 experiments.

| | **Biotin-CGRP(8–37)** | **Biotin-ssCGRP(8–37)** |
|---|---|---|
| $k_{on}$ ($M^{-1}s^{-1} \pm$ SEM) | $6.33 \times 10^4 \pm 4.77 \times 10^3$ | $2.90 \times 10^5 \pm 5.24 \times 10^3$ |
| $k_{off}$ ($s^{-1} \pm$ SEM) | $3.31 \pm 0.31$ | $0.0077 \pm 0.0003$ |
| $K_D$ (nM $\pm$ SEM) | $52{,}200 \pm 1{,}266$ | $26.4 \pm 1.1$ |
| Residence time[1] (s) | $0.31 \pm 0.028$ | $131.2 \pm 5.6$ |
| $t_{1/2}$[2] (s) | $0.21 \pm 0.02$ | $90.9 \pm 3.9$ |
